# Geminiviral βC1 orchestrates organellar genomic instability to augment viral infection by hijacking host RecA

**DOI:** 10.1101/2021.12.29.474444

**Authors:** Ashwin Nair, C.Y. Harshith, N. Anushree, P. V. Shivaprasad

## Abstract

Chloroplast is the site for transforming light energy to chemical energy. It also acts as a production unit for a variety of defense-related molecules. These defense moieties are necessary to mount a successful counter defence against pathogens including viruses. Geminiviruses disrupt chloroplast homeostasis as a basic strategy for their successful infection inducing vein-clearing, mosaics and chlorosis in infected plants. Here we show that a geminiviral pathogenicity determinant protein βC1 directly interferes with plastid homeostasis. βC1 was capable of inducing organelle-specific nuclease to degrade plastid genome as well as diverted functions of RecA1 protein, a major player in plastid genome maintenance. βC1 interacted with RecA1 in plants and its homolog in bacteria to reduce the ability of host cells to maintain genomic integrity under stresses. Further, reduction in the coding capacity of plastids severely affected retrograde signalling necessary for viral perception and activation of defense. Induction of chloroplast-specific nuclease by βC1 is similar to phosphate starvation-response in which nucleotides are recycled to augment synthesis of new, potentially viral, DNA. These results indicate the presence of a novel strategy in which a viral protein alters host defence by targeting regulators of chloroplast DNA. We predict that the mechanism identified here might have similarities in other plant-pathogen interactions.

**One line summary:** βC1 alters plastid genome metabolism.

## Introduction

Chloroplast is an emerging hub for defense signalling during plant-pathogen interactions (de Torres Zabala et al., 2015; Padmanabhan and Dinesh-Kumar, 2010; Nomura et al., 2012; Serrano et al., 2016). In addition to being at the centre for photosynthesis and various metabolic processes, chloroplast also synthesizes various immune modulators such as salicylic acid (SA), jasmonic acid (JA), ethylene (ET), abscisic acid (ABA), various secondary metabolites, aromatic amino acids and other signalling molecules such as H_2_O_2_, ROS, and singlet oxygen species (^1^O_2_) (Wildermuth et al., 2001; León and Sánchez-Serrano, 1999; Nambara and Marion-Poll, 2005; Chan et al., 2010). The production of these immune modulators is tightly regulated to avoid dysfunctional expression leading to negative growth effects (Chandran et al., 2014).

Plants being sessile are continuously threatened by biotic and abiotic stresses. They have evolved an intricate signalling network for recognition of pathogens. Pathogenic markers (PAMPs, DAMPs, MAMPs, etc.) once recognised by PATHOGEN RECOGNITION RECEPTORS (PRR) on the cell surface or in the cytoplasm, activate signalling to the nucleus via MAPK pathway, and to other organelles such as chloroplast, peroxisomes and mitochondria via an unknown pathway (Choi and Klessig, 2016; Ma et al., 2016; Grant and Jones, 2009; Nomura et al., 2012). In response to pathogen-derived signal, chloroplast generates ROS, inducing retrograde signalling with the nucleus leading to a transcriptional induction, synthesising various defense-related genes and producing hormones such as SA in the plastid stroma (Chan et al., 2016; Grant and Jones, 2009). SA synthesis in turn induces the expression of defense genes responsible for pattern-triggered immunity (PTI) and effector-triggered immunity (ETI), thus limiting pathogenic spread (Seyfferth and Tsuda, 2014; Pieterse et al., 2012). Activation of PTI is also a part of the antiviral arsenal in plants (Machado et al., 2015; Kørner et al., 2013; Iriti and Varoni, 2015; Nicaise and Candresse, 2017).

Chloroplast consumes significant cellular resources. For accommodating the translational load, plastid genomes are present in multiple copies, and despite being relatively smaller, they account for substantial DNA content of the cell (>20% in mature leaf) (Rauwolf et al., 2010; Sakamoto and Takami, 2018). Multiple copies of the plastid genome are essential for maintaining homeostasis during various metabolic processes (Bendich, 1987; Udy et al., 2012). The chloroplastic genome is maintained by poorly studied organelle-specific DNA DAMAGE AND REPAIR (DDR) machinery. Orthologs of bacterial RecA proteins like RecA1, RecA2, DRT100 and DRT102, along with repair proteins like MUTATOR S (MutS) are members of DDR family and they play an essential role in maintaining the copy number and structure of chloroplast DNA (cpDNA) (Majeran et al., 2012; Rowan et al., 2010; Odahara et al., 2017).

Most DNA viruses accumulate in the host nucleus and depend on their host for replication (Schmid et al., 2014). Among the viruses that infect plants, geminiviruses are the largest family of single-stranded DNA (ssDNA) viruses. Geminiviral particles are directly injected into the phloem by insect vectors surpassing the primary layer of defence in plants (Hanley-Bowdoin et al., 2013; Rizvi et al., 2015). However, recent studies indicated the activation of innate immunity upon geminiviral infection. The wounding response triggered by insect vector feeding can prime PTI and RNA-interference (Wang et al., 2021). Similarly, DAMPs and PAMPs are recognised by cell surface receptors such as RECEPTOR LIKE KINASES (RLKs) and RECEPTOR LIKE PROTEINS (RLPs) (Teixeira et al., 2019; Niehl et al., 2016; Nicaise and Candresse, 2017; Zorzatto et al., 2015). In line with the role of chloroplast in antiviral defense, various viral effectors disrupt the key process of chloroplast metabolism to sabotage PTI activation (Fondong et al., 2007; Gnanasekaran et al., 2019; Bhattacharyya et al., 2015; Nair et al., 2020; Medina-Puche et al., 2020).

Geminiviruses employ robust mechanism for replication, involving both rolling-circle replication (RCR) and recombination-dependent replication (RDR) strategies. RCR is a robust process but also leads to the production of heterogeneous ssDNA of varying lengths due to polymerase runoff or improper termination (Heyraud et al., 1993; Heyraud-Nitschke et al., 1995; Stanley, 1995). Evidence suggests the role of DNA repair machinery in the repair and reconstruction of intermediate replicative forms (RFs) (Jeske et al., 2001; Ascencio-Ibáñez et al., 2008; Preiss and Jeske, 2003). RecA and Rad proteins are the mediators of homologous recombination and act as core proteins of DDR and SOS-repair pathways in organisms (Maslowska et al., 2019; Chappell et al., 2016). There are also a few studies of geminiviral proteins influencing host DDR machinery. Geminiviral Rep interacts specifically with Rad54, an important player in homologous recombination (Kaliappan et al., 2012). Choudhury et al. (2013) observed induction of Rad51 transcription in MYMV-infected plants, suggesting possible exploitation of DNA repair genes by the virus (Suyal et al., 2013). Rad51D appeared to act as an important player in maintaining the genomic integrity of early viral replication intermediates (Richter et al., 2016). Surprisingly, *Radish leaf curl virus* (RaLCV) βC1 protein has an ATPase activity but the relevance of this activity in connection to DDR or cell-cycle pathways is unknown (Clerot and Bernardi, 2006; Gnanasekaran et al., 2021).

The expression of chloroplast-localized βC1, a geminiviral pathogenicity determinant, is toxic to plants (Cui et al., 2004; Yang et al., 2008). We and others had previously reported a multitude of growth defects observed in host plants upon ectopic expression of βC1 (Nair et al., 2020; Bhattacharyya et al., 2015; Cheng et al., 2011; Briddon et al., 2003). Here we show that *Synedrella yellow vein clearing virus* (SyYVCV) βC1 (Das et al., 2018) induces selective degradation of cpDNA during viral infection. βC1 achieved this by inducing expression of DPD1 nuclease. βC1 was also able to interact and modulate the function of RecA1, a chloroplastic DDR protein in plants and its ortholog RecA in bacteria. Interaction of βC1 with RecA1 was paramount for successful viral pathogenesis, in increasing the viral titre and for the formation of symptoms. Disruption of chloroplastic homeostasis by degradation of cpDNA interrupted PTI signalling, further curtailing SA synthesis and SYSTEMIC ACQUIRED RESISTANCE (SAR). We show that βC1 can induce genotoxic stress in plants and alter the expression of various important DDR pathway genes. Our results indicate that interaction of βC1 protein with a DDR protein RecA1 and its influence on genotoxicity is another novel aspect of the much-appreciated arms race between viruses and their host plants.

## Results

### βC1 alters the expression of key regulatory genes in host plants

The βC1-expressing transgenic tobacco plants were sterile, stunted, chlorotic, and presented an early flowering phenotype with exerted stigma (Figure 1A and 1B). As observed previously, the toxicity of the βC1 protein was dampened upon C-terminal tagging (Figure S1A). To understand the cellular pathways affected by βC1, we performed a transcriptome analysis using the Illumina Hi-Seq platform. We obtained an average of 20 million (M) x 2 paired-end reads, out of which 92% matched to *N. tabacum* genome. Upon further characterization of 3576 genes showing maximum differential expression, 1963 genes were found up-regulated and 1613 genes were down-regulated (Figure S1B and S1C). Various defense response regulators were found misexpressed as expected for a pathogenicity determinant protein like βC1. Innate immune regulators such as secondary metabolite (suberin, lipids, and phenylpropanoid) pathway genes were down-regulated in βC1 transgenic plants (Figure S1D). We hypothesized that most of the phenotypes observed in βC1 transgenic plants might be due to disrupted signalling pathways, prominently hormone and circadian rhythm pathways responsible for maintenance of development and vegetative to flowering transition, respectively. In agreement with this, transcripts of GIGANTEA (GI-like) and CONSTANS (CO5-like) key regulators of circadian rhythm were 7 and 3.5-fold upregulated, respectively, in βC1 transgenic plants. Chlorophyll A/B binding proteins which are under the control of TIMING OF CHL A/B EXPRESSION1 (TOC1) were 9.5-fold upregulated. A regulator of flowering LATE ELONGATED HYPOCOTYL (LHY) homolog was 6-fold down-regulated, while its counterpart EARLY FLOWERING 4 (ELF4) was 8-fold up-regulated. As observed previously (Zhao et al., 2021), PHYTOCHROME-INTERACTING FACTOR4 (PIF4), a secondary metabolism regulator was 5.4-fold upregulated. In addition, multiple auxin-responsive proteins including YUCA11 were down-regulated in these plants, while cytokinin degrading enzyme CYTOKININ DEHYDROGENASE 7-LIKE was 4-fold up-regulated (Figure S1E) suggesting deregulation of hormonal signalling in these plants.

**Figure 1:**
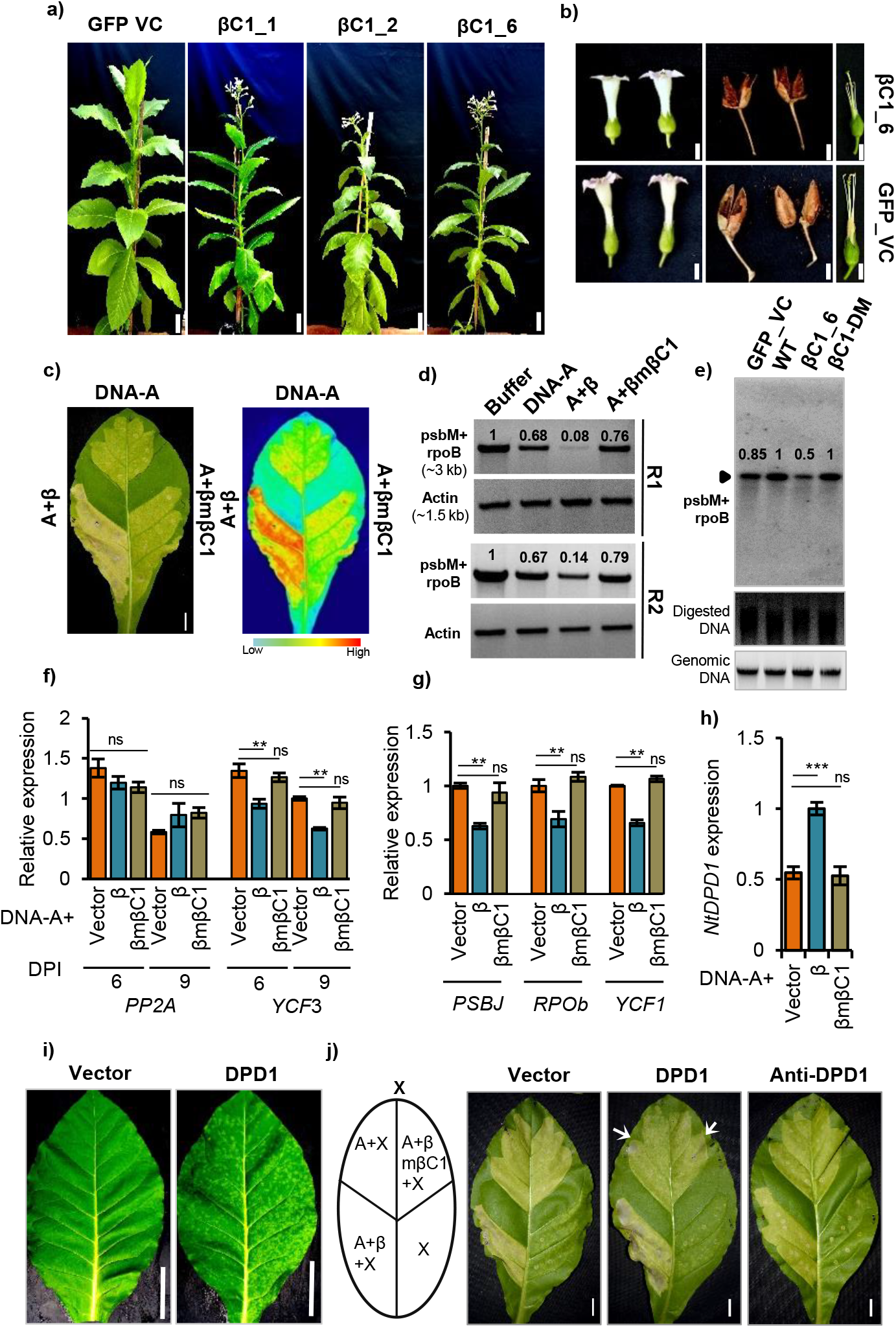
βC1 selectively degrades chloroplastic DNA. **a)** Phenotype of transgenic *N. tabacum* lines over-expressing βC1. Pictures taken at 50 days post transplantation (DPT). N=3 for each transgenic lines **b)** Pictures of flower and seedpod. **c)** *N. tabacum* leaves showing chlorosis when infected with DNA-A, DNA-A+β and DNA-A+βmβC1 (left). Heat map representing the extent of chlorosis (right) **d)** Semi-quantitative DNA PCR showing abundance and integrity of chloroplastic (psbM+rpoB) or nuclear (actin) genome upon infection with β or βmβC1. **e)** Southern blot (SB) showing the abundance of chloroplastic genome in βC1 OE plants. A 3-kb (psbM+rpoB) chloroplastic region was used as probe. Genomic DNA panel represents a duplicate gel with same amount of DNA used in SB for normalization. **f)** DNA qPCR representing the abundance of chloroplastic (*YCF3*) or nuclear (*PP2A*) genes upon infection with β or βmβC1. Nuclear *ACTIN* was used for normalization. **g)** Same as f) except 9 DPI sample, with other plastid genes. **h)** Expression of *DPD1* nuclease in virus infected samples with βC1 and mβC1. **i)** Phenotype of the systemic leaf expressing *NtDPD1* (PVX-*NtDPD1*). 25 DPI. N=4, image linked with figure S2. **j)** Chlorosis and necrotic phenotypes of *N. tabacum* leaves infected with DNA-A, DNA-A+β or βmβC1 co-infiltrated with PVX vector (left panel), P-*DPD1* (middle panel) or P-*anti-DPD1* (right panel). White arrow highlights new necrotic spots. Biologically replicated twice. Scale bar 2 cm. N-terminal GFP tagged βC1 was used for making transgenic βC1 lines. Size bar in a) and b) corresponds to 5.8, and 1 cm. Size bar in c), i) and j) is 1.2 cm. GFP VC is GFP over-expressing vector control plant. Tukey’s multiple comparison test with three stars representing P-value, P ≤ 0.001 and two stars P ≤ 0.01. n=4.

Surprisingly, we also observed differential expression of a set of DDR genes involved in genome maintenance and repair in βC1-expressing transgenic plants (Figure S2A). Although previous research suggested deregulation of DDR genes during geminiviral replication (Ascencio-Ibáñez et al., 2008), it was not known which viral protein was involved. βC1 plants showed up-regulation of various DDR genes such as Rad50-like, photolyases, DRT proteins, BRCA1-like, J-Domain proteins, several nucleases and helicases (Figure S2A). Majority of the DDR genes that were upregulated in βC1 plants were necessary for the maintenance of the chloroplast genome (Day and Madesis, 2007). Proteins such as DRT102, DRT100, DPD1 nuclease, ARC6, and FtsZ proteins regulate the copy number, replication, and damage repair of the plastid genome (Figure S2A). We also observed deregulation of many chloroplast localized genes in βC1 transgenic plants (Figure S2B and C), correlating with the symptoms observed in these plants. Similar deregulation of plastid genes was observed in chloroplast-localized RALCV βC1 (Bhattacharyya et al., 2015), where chloroplastic ultrastructure was compromised leading to the loss of photosynthetic output.

### βC1 induces chlorosis by destabilising the plastid genome

Based on phenotype as well as changes in host DDR genes, we hypothesised that the cause for the deregulation of such a huge number of chloroplast genes is likely due to selective plastid DNA destabilization by βC1. Chloroplast localized SyYVCV βC1 was previously identified as the causal protein for the symptoms during viral infection (Nair et al., 2020). To further confirm that βC1 is the protein responsible to induce chlorosis during viral infection, we infected *N. tabacum* leaves with SyYVCV DNA-A alone or with DNA-β and DNA-β with point mutations in βC1 ORF (DNA-βmβC1 (mSIM2,3,4;,(Nair et al., 2020)). The point mutations in the SIM region of βC1 completely abolishes βC1 functions. Strong chlorosis was observed in the segment of leaves infected with DNA-A+β when compared to DNA-A alone. Further, very mild chlorosis similar to DNA-A was observed in the segment infected with DNA-A+βmβC1 (Figure 1C). To check if the stability of the plastid genome was compromised in these leaves, we analysed abundance and integrity of plastid genome by amplifying a 3-kb segment (psbM and rpoB) from the plastid genome. A drastic reduction in plastid DNA was observed in the presence of DNA-A+β as compared to control nuclear DNA fragment (Figure 1D). Further, we analyzed the abundance of plastid DNA in βC1 transgenic plants *via* Southern blotting (SB) analysis. A clear reduction in the plastid DNA in βC1 transgenic plants as compared to vector control or functionally inactive βC1-DM (βC1, C-terminal tagged) plants was observed (Figure 1E). The instability and degradation of cpDNA caused by βC1 was more evident in later stages of infection. These observations were further validated with other plastid genes using qPCR (Figure 1F and G). *PP2A* and *ACTIN* gene taken as a proxy for nuclear genome was relatively stable in different samples as compared to the chloroplastic markers (*YCF3*, *YCF1*, *PSBj* and *RPOb*). Previous studies had highlighted a conditional role for plastid nuclease DPD1 in degrading plastid genome (Sakamoto and Takami, 2018). *DPD1* was 4-fold upregulated in βC1 transgenic plants (Figure S2A). We analysed the expression of *DPD1* during viral infection and observed a significant induction in the presence of DNA-β, but not in DNA-βmβC1 (Figure 1H). We explored if the cause for βC1-induced chlorosis and necrosis during infection is linked to DPD1. Chlorotic and necrotic mosaic patches were observed upon over-expression of DPD1 (PVX-*NtDPD1*) in *N. tabacum* and *N. benthamiana* plants (Figure 1I and Figure S2D and E). To further explore the role of *NtDPD1* in βC1 induced chlorosis during viral infection, *DPD1* or antisense-*DPD1* were expressed along with DNA-A, DNA-A+β or DNA-βmβC1 in *N. tabacum* leaves (Figure 1J). As expected, WT βC1 alone induced chlorosis and limited necrosis in vector control leaves (Figure1J, left panel). The necrosis was significantly enhanced in *DPD1* over-expressing leaves if infiltrated along with DNA-β (Figure1J, middle panel). Necrosis or chlorosis was not observed in DNA-β or DNA-βmβC1 in antisense-*DPD1* expressing leaves (Figure1J, right panel). These results suggested that plastid DNA was selectively destabilised by βC1-induced DPD1 during viral infection.

### βC1 induces genotoxicity in bacteria

To gain further insight into the plastid DNA destabilization by βC1, we devised a bacterial cell-based genotoxicity assay. Since geminiviruses have bacterial origin and chloroplast is also derived from bacteria (Krupovic et al., 2009), we hypothesized that the expression of βC1 might be deleterious in bacteria via similar mechanism observed in chloroplast. βC1 and its N and C-terminal truncation mutants were induced in Rosetta-Gammi (DE3) cells followed by a treatment of a sub-lethal dose of UV-C or bleomycin to induce DNA damage (Figure 2A). The UV-C dose had minimal to no effect on the viability of cells with active DDR machinery, such as in DE3 protein expressing cells. The MBP control protein expressing cells showed appropriate growth before and after induction following stress, suggesting the external sub-lethal DNA damage was sustained and repaired in these cells. As expected, βC1 expressing cells showed acute lethality, suggesting that βC1 is genotoxic to bacteria, similar to plants. Induction of βC1 minimally reduced the viability of cells as seen in the drop assay, while the addition of a sub-lethal dose of UV tipped the balance between DNA damage and repair, towards damage and cell lethality. Interestingly, βC1 δC59 expressing cells did not show lethality upon induction in UV or bleomycin, whereas βC1 δN59 expressing cells showed cell death similar to full-length βC1. These results suggest that the plastid DNA genotoxicity inducing mechanism of βC1 is also conserved in bacteria, but in bacteria the βC1 induced genotoxicity is conditional suggesting a role of DDR repair machinery.

**Figure 2:**
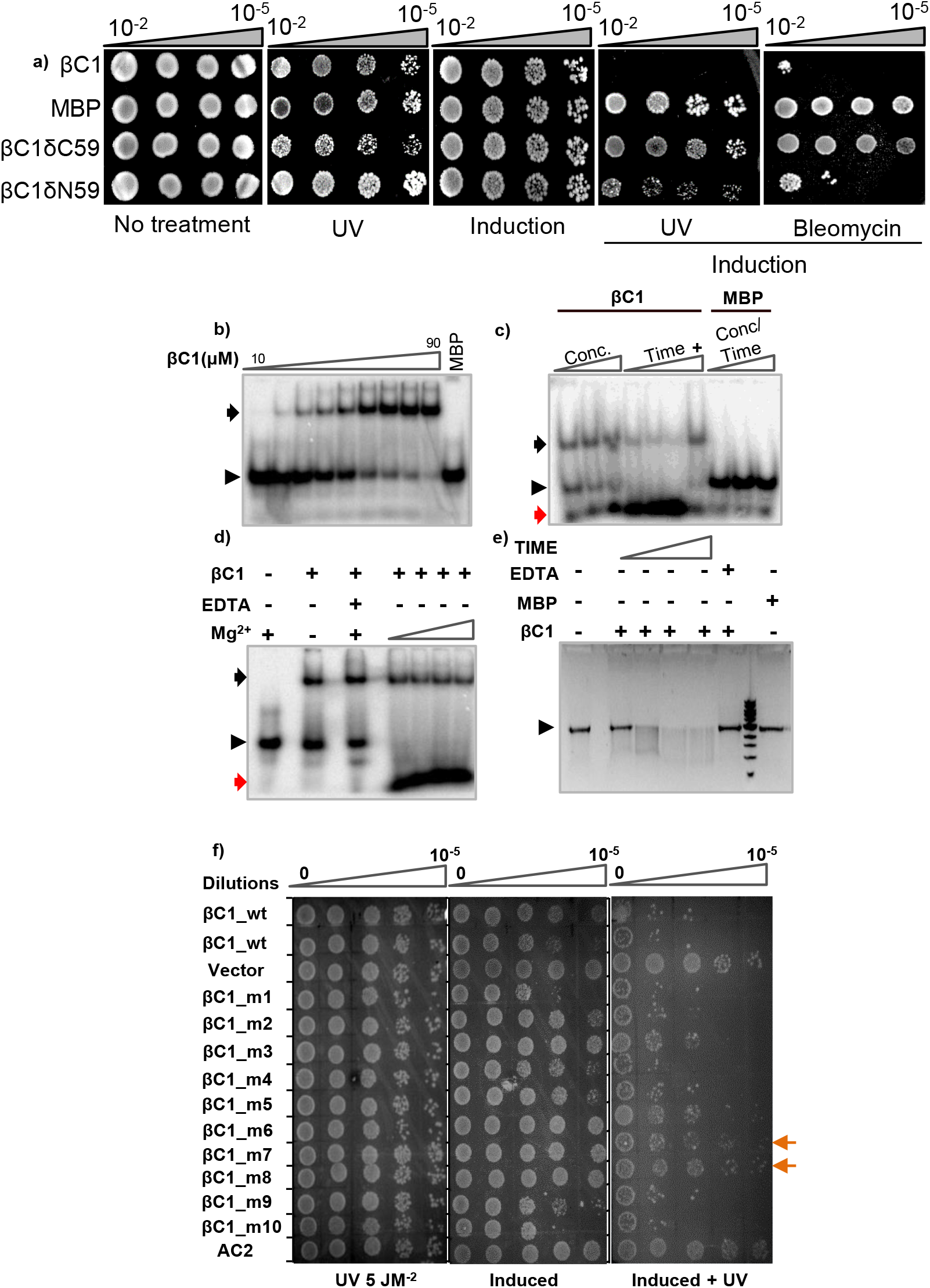
SyYVCV βC1 displays genotoxicity in *E.coli* and associated nuclease activity *in-vitro*. **a)** DNA damage sensitivity assay: βC1 was transformed in Rosetta Gammi DE3 cells and grown till mid lag phase followed by spotting on LB agar with sub-lethal UV: 5 J/m2, 254 nm. Bleomycin 1 µg/ml. **b)** EMSA showing binding of βC1 with ssDNA in a 8% native PAGE gel. Triangle indicates increasing concentration of βC1. **c) Nuclease assay:** βC1 was incubated with ssDNA in various concentrations for varying duration followed by EMSA in a 6% native PAGE gel. (+) indicates addition of 1 mM EDTA. **d) Cation dependency assay:** Mg^2+^ (1 to 5 mM) or EDTA (5 mM) was co-incubated with βC1 and ssDNA followed by EMSA on a 6% native PAGE gel. **e) Endonuclease activity assay:** βC1 was incubated (1 to 4 h) with covalently closed circular ssDNA (ɸ174) followed by visualization on 1% agarose gel. **f) Genotoxicity assay:** βC1 and mutants were spotted on plates in various dilutions and treatments. Induced represents induction of gene with 0.1 mM IPTG, UV is UV-C (254 nm). All assay results were replicated multiple times. Vector expresses MBP. Additional info: N-terminal MBP tagged βC1 (*E. coli* purified, SEC, DEAE) was used in all biochemical assays. MBP parallelly processed with βC1 was used as control. ssDNA substrate used was 49-nt long (150 pg). ssDNA was labelled at 5’ using ^32^P. Black and red arrows indicate bound and degraded fraction of ssDNA, respectively. Black triangle-unbound substrate.

To understand the biochemical mechanism behind the genotoxicity of βC1, we recombinantly expressed and purified βC1 from *E. coli* using size-exclusion and ion-exchange chromatography. As βC1 from other viruses have been shown to bind to different nucleic acids, and chloroplastic DNA was targeted by βC1 in SyYVCV infected plants, we explored if SyYVCV βC1 can bind ssDNA and/or double-stranded (ds) DNA and RNA substrates and if it can alter stability of nucleic acids. The SyYVCV βC1 was able to bind both ssDNA (Figure 2B) and dsDNA (Figure S3A), but displayed significantly higher binding to ssDNA (Figure 2B). The strength of binding to dsDNA was directly proportional to its length (Figure S3A). However, βC1 did not exhibit significant binding to either ss or dsRNA substrates (Figure S3B to E). We observed significant degradation of ssDNA *in vitro* in the presence of βC1 in multiple biological replicates. The control MBP protein showed neither binding nor nuclease activity *in vitro* (Figure 2C). Since nucleases mostly require divalent cations as a cofactor, we checked metal ion dependency for the associated nuclease activity of βC1 and observed optimum degradation of ssDNA in presence of Mg^2+^ ions and to a lesser extent with Mn^2+^ (Figure S3F). We next validated these results using a metal ion chelator EDTA that removes Mg^2+^ from the catalytic interface and did not observe any nuclease activity (Figure 2D and S3G). Interestingly, the ability of βC1 to bind ssDNA was not compromised in the presence of EDTA, suggesting an Mg^2+^ independent binding (Figure 2D). Incubation of purified βC1 with circular ssDNA led to its complete degradation, suggesting that the *in vitro* nuclease activity associated with βC1 has both endonuclease as well as exonuclease activity (Figure 2E). Similar results were observed upon incubation of βC1 with plasmid DNA (Figure S3H). Interestingly, plant DPD1 is also an endonuclease and exonuclease degrading both ss and dsDNA. Even though, studies suggest DPD1 is not of endosymbiotic origin, it is likely that βC1 might be regulating other structurally conserved exonuclease family members in bacteria (Takami et al., 2018; Sakamoto and Takami, 2018). Combining all these observations we conclude that *in vitro* purified βC1 has a novel associated nuclease activity that might be involved in cellular genotoxicity.

### Specific domains of βC1 mediate genotoxicity, multimerization and DNA binding properties

To delineate the motif associated with nuclease activity, we made multiple point mutations (mutant 1 to 10) in βC1 (Figure S3I). These point mutations were made based on their conservation across different βC1 sequences derived from different viruses. All mutants were recombinantly purified similar to WT βC1 (Figure S3J), and their DNA binding ability was analysed along with WT βC1 as a positive control. MBP acted as a negative control. While controls behaved as expected, mutants 6 and 8 showed significant reduction in binding to ssDNA (Figure S3K). None of these point mutations completely abolished the observed nuclease activity. However, we observed a reduction in the nuclease activity of mutant 6, 7 and 8 (Figure S3L).

To further delineate the genotoxicity domain of βC1, we used cell-based UV-genotoxicity assay. βC1 and its point mutants were induced in Rosetta-Gammi (DE3) cells followed by a treatment of a sub-lethal dose of UV-C (Figure 2F). We used MBP, and SyYVCV AC2, another nucleic acid-binding protein of SyYVCV that lacks associated nuclease activity, as controls. As previously observed, MBP and AC2 expressing cells showed appropriate growth before and after induction following stress, suggesting the role of proactive DDR machinery. As expected, βC1 expressing cells showed acute lethality after sub-lethal UV stress. A few βC1 mutants (mutants 10 and 1) showed enhanced lethality during induction as well as with UV-stress when compared to WT βC1. Fascinatingly, mutants 7, 8 and 6 showed a significant decrease in cell lethality (Figure 2F). These results suggested that βC1-associated nuclease activity was capable of causing genotoxicity in cells and its C-terminus domain is involved in this activity.

βC1 exists as multimers *in vivo* as well as in purified fractions (Cheng et al., 2011). Multimerization might influence the DNA binding of proteins. We recombinantly expressed and purified βC1 from *E. coli* and observed higher-order multimers as well as monomers (Figure 3A). To delineate the multimerization motif, we made various truncation mutants of βC1. WT βC1 protein eluted peak was at 43 ml, suggesting that βC1 protein (MBP-βC1, ∼59 kDa) formed multimers in solution (Figure 3A). All the truncation mutants of βC1 except δN59 were able to form multimers. Careful examination of the truncation mutants, fine mapped the minimum multimerization motif between residues 51 to 59 (Figure 3B and S4A to L). The δC59 showed multimerization similar to WT βC1, whereas in δN59, the multimerization activity was completely abolished (Figure 3C). Interestingly, the DNA binding ability of the δN59 mutant was significantly reduced when compared to other mutants matching the requirements for multimerization (Figure 3D and E), suggesting that multimerization and DNA binding activities are linked together. Interestingly, the C-terminus is essential for observed genotoxicity in bacteria (Figure 2A), but the DNA binding motif is mostly localised in the N-terminus (Figure 3D). These results suggests that βC1 C-terminus can indirectly induce genotoxicity, likely by inducing nuclease.

**Figure 3:**
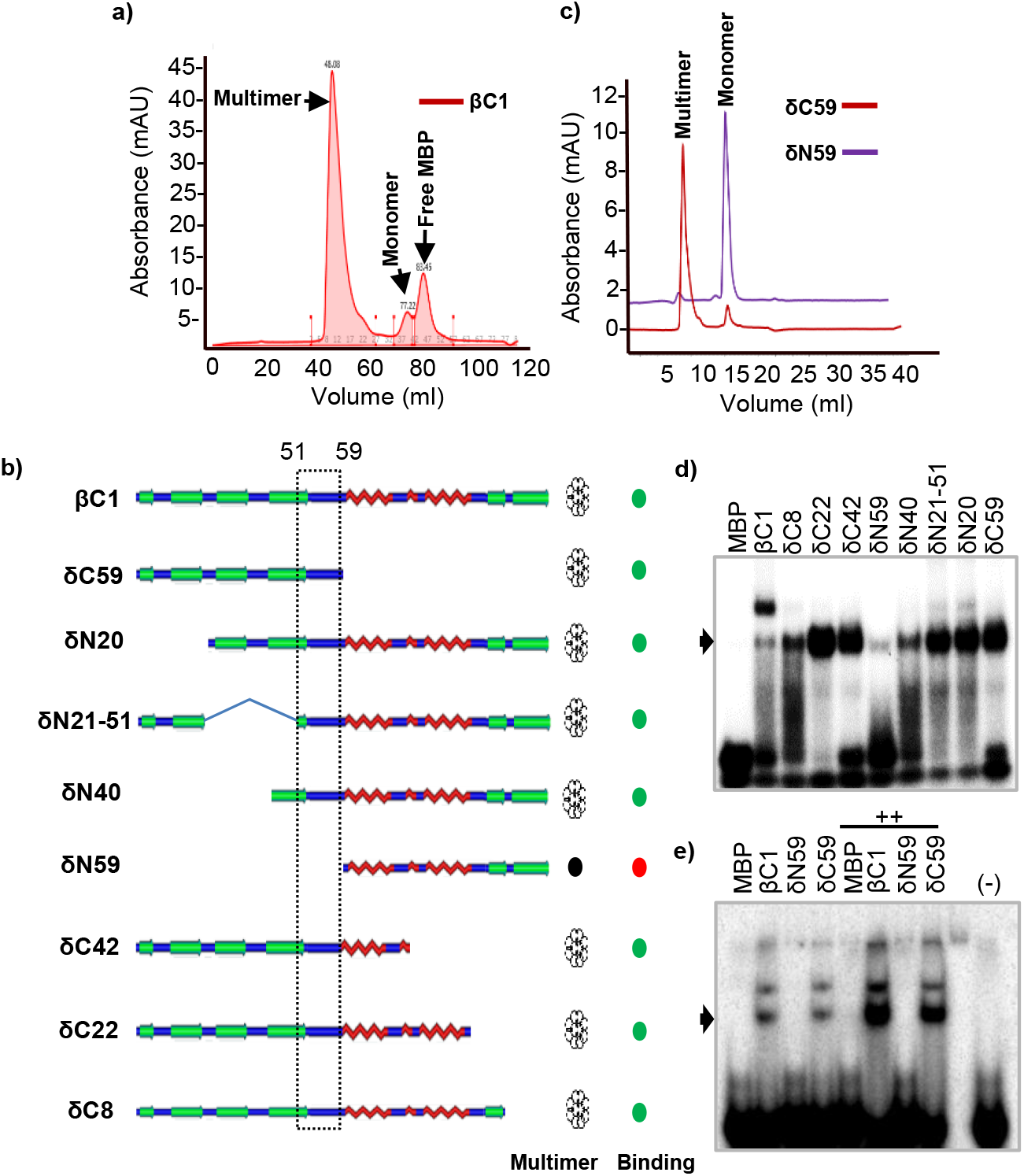
Demarcation of multimerization domain of βC1. **a)** Size exclusion profile of MBP-βC1 on a SD200 preparative column highlighting multimer, monomer and free MBP in βC1 fractions. **b)** Schematics of βC1 truncation mutants showing a summary of size exclusion analysis in a SD200 sephadex analytical column. Right side columns depict multimerization and DNA binding state of respective protein. Green and red circles indicate presence or absence of DNA binding, respectively. **c)** Overlaid size exclusion profile of C-terminal (δN59) and N-terminal (δC59) truncation mutants of βC1 analysed on an SD200 analytical column. **d)** EMSA showing ssDNA binding of βC1 and its truncation mutants. **e)** Same as d) except N and C terminal truncation mutants were used. ++ indicates 2-fold increase in protein and (-) indicates no protein control. Black arrow indicates binding.

### βC1 expressing bacterial cells require RecA for survival

*In vitro* DNA binding assays, purified βC1 had associated nuclease activity but in bacterial cells genotoxicity was conditional. Bacteria deals with DNA damaging stress employing an evolutionary conserved family of proteins such as RecA recombination protein. RecA family of proteins maintain the integrity of bacterial genome and its homologs are also conserved in chloroplast genome (Maslowska et al., 2019; Rowan et al., 2010). In bacterial cells, RecA protein acts as the central regulator of DDR machinery. We hypothesized a direct role of RecA in DNA damage response induced by βC1, since this group of genes were more responsive in transcriptome analysis. We used BLR (DE3) cells that lack a functional RecA protein in genotoxic assay. MBP control or empty vector did not show any difference in growth among treatments, however, as expected, βC1 expressing cells did not survive the induction in BLR (DE3) (Figure S5A and B). We tested the effect of βC1 mutants in BLR cells and observed that mutant 7, 8 and mβC1 rescued cells from lethality, reinforcing results observed in UV assay, and highlighting role of C-terminal in genotoxicity (Figure S5C and Figure 2F). To verify RecA as a key player, we complemented *Caulobacter vibrioides RecA* (Cv*RecA*) in βC1 expressing BLR (DE3) cells. The *CvRecA* protein is a close homolog of *E. coli RecA* (*EcRecA*), and this complementation completely abrogated βC1-induced genotoxicity (Figure S5D). These results suggest that RecA is required by bacterial cells to subdue βC1-induced genotoxicity.

We used truncation mutants of βC1 to delineate the minimum motif required for genotoxic effects in bacteria by employing both UV-stress and BLR (DE3) based assays. MBP was used as control. As expected, MBP, did not show any lethality in the UV and BLR assays, whereas βC1 expressing cells presented acute lethality. Interestingly, truncating even as minimum as 8 residues (δC8) from the C-terminus significantly altered the genotoxic activity of βC1 (Figure 4A). Careful examination of the results showed a complete loss of lethality upon truncating 42 residues (δC42) from the C-terminal. Surprisingly, we observed an increase in genotoxicity in N-terminal truncation mutants of βC1. δN20 and δN21-41 showed increased genotoxicity as compared to WT βC1 (Figure 4A to C). Similar results were also observed in point mutants of βC1 (Mutant 10) that has substitutions in the N-terminal 20 residues (Figure 2F). These results suggested that the C-terminal of βC1 induces genotoxicity in cells while its N-terminus might be essential for regulating this activity.

**Figure 4:**
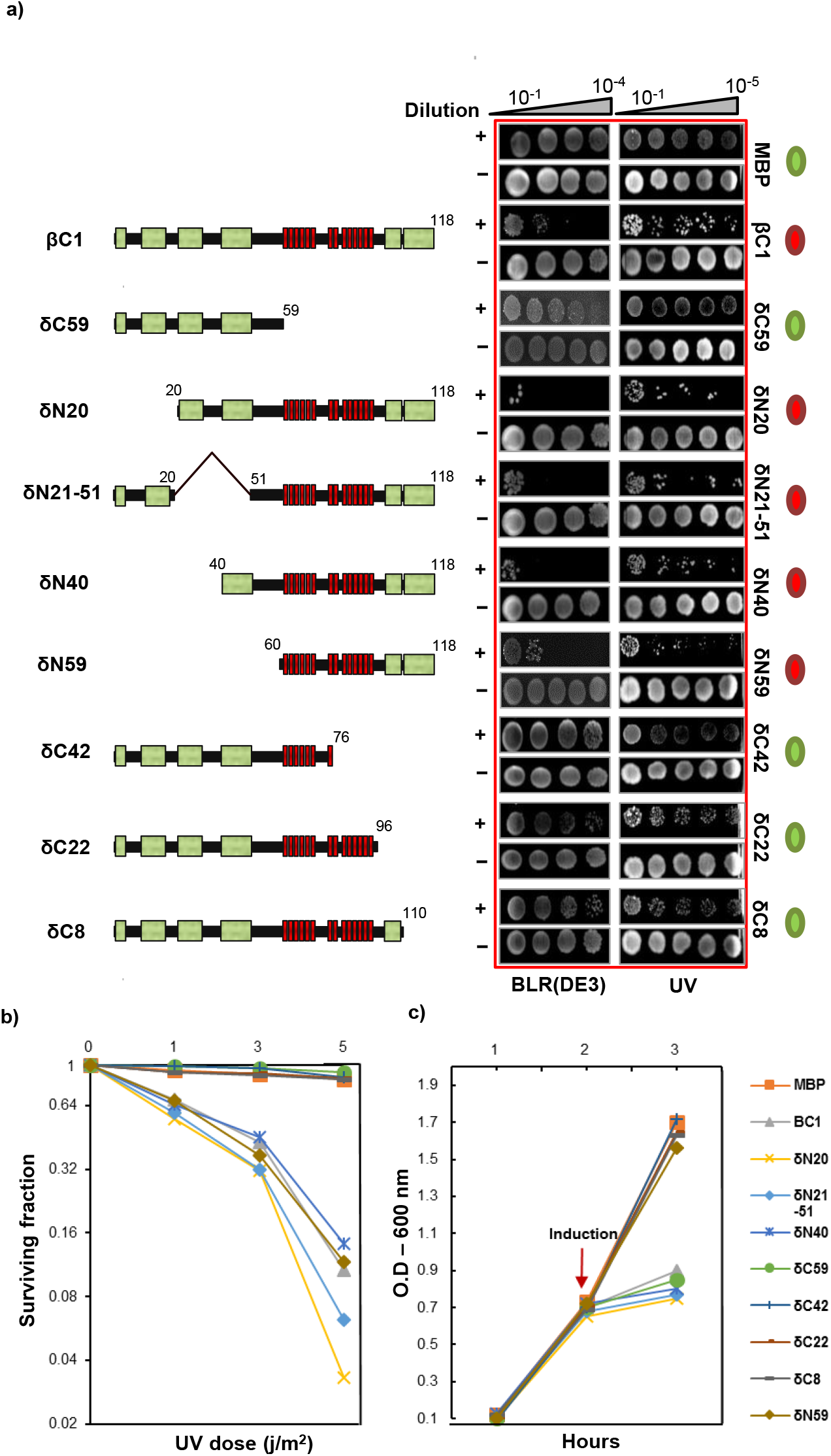
SyYVCV βC1 C-terminal end is required for genotoxicity. **a)** Schematic diagram of βC1 mutants (left panel). βC1 and its mutants were transformed in BLR (DE3) cells and induced with IPTG (middle panel). Right panel: βC1 was expressed in Rosetta Gammi cells followed by spotting and exposure to UV-C. **b) and c)** Plots showing survival rate and growth after c) UV treatment and d) induction of βC1 in BLR cells (0.2 mM). Green and red circles represent majority fraction of cells being alive or dead.

### Bacterial RecA physically interacts with geminiviral βC1

DNA repair response or SOS repair in bacteria is a multi-component system, and RecA protein acts as a central player (Maslowska et al., 2019). It was previously observed that virulence proteins such as the pathogenicity determinants (for example Rep protein) interacted with Rad proteins. Geminiviral Rep protein interacts with *S. cerevisiae* Rad54 (Kaliappan et al., 2012). βC1 BLR (DE3) genotoxicity assay further hinted to a direct role of RecA in modulating activity of βC1. We explored if βC1 is capable of interacting with RecA. Using RecA specific antibody, we were able to detect RecA in purified βC1 fractions. A 37-kDa band was detected in the purified βC1 protein fraction but not in the MBP fraction. Interestingly, RecA was detected in the C-terminus truncation mutant (δC59) of βC1 but not in the N-terminus truncation mutant (Figure 5A).

**Figure 5:**
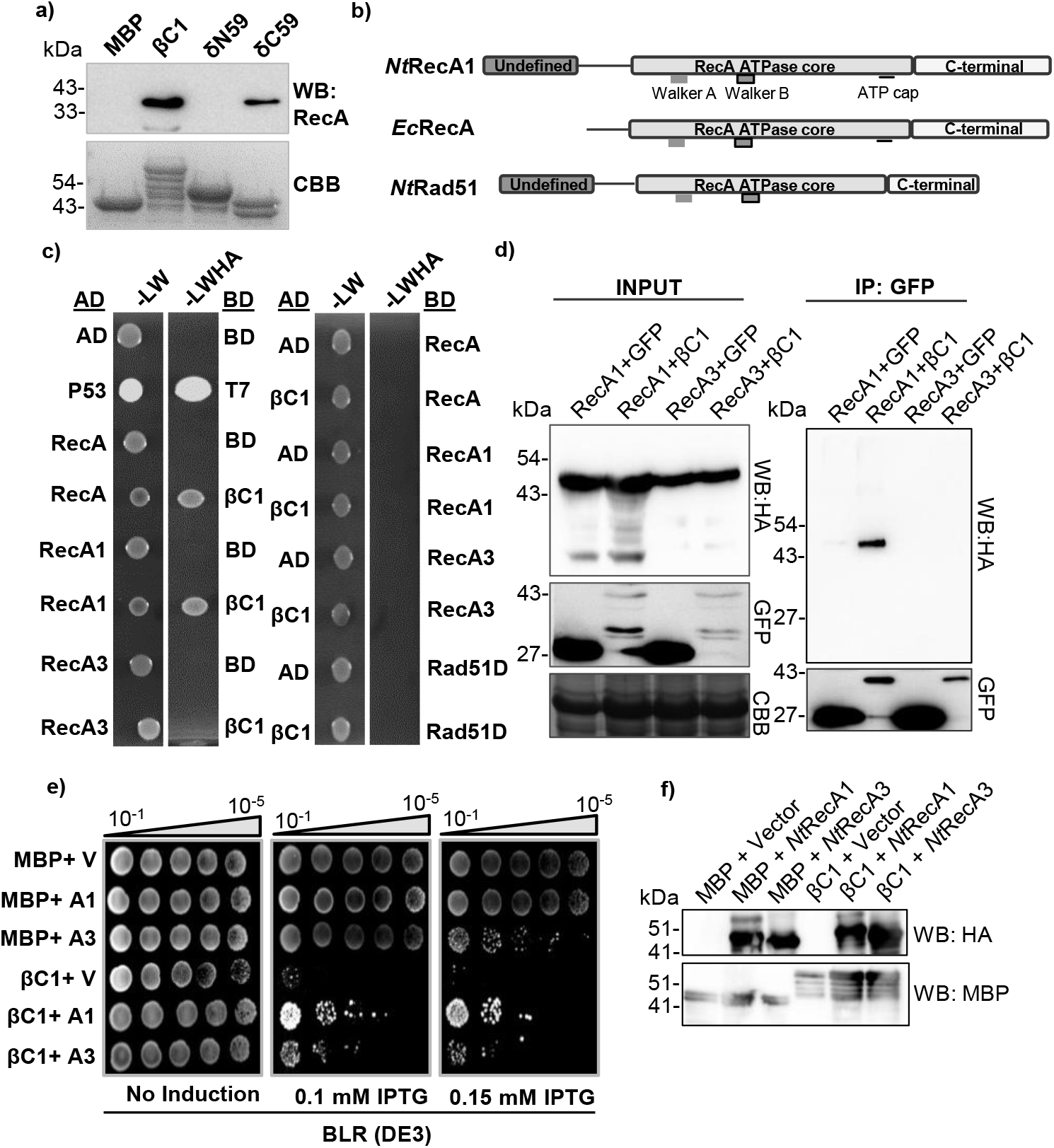
*Nt*RecA1 interacts with SyYVCV βC1. **a)** WB of recombinantly purified βC1 with anti-RecA showing the presence of *Ec*RecA in purified βC1 fractions, n=3. **b)** Domain architecture of RecA and its plant homologs. **c)** Y2H assay showing interaction of βC1 with RecA and its plant homologs. Dilution showed is 10^-2^. The assay was performed with different dilutions and knockout media, representative image shown. **d)** *in planta* pull-down assay to check interaction of βC1 with RecA homologs. RecA homologs were tagged with HA and βC1 was tagged with GFP. GFP was taken as control. **e)** Complementation of *Nt*RecA1 and *Nt*RecA3 in MBP or βC1 expressing BLR (DE3) (recA^-^) cells. **f)** WB confirmation of complementation experiment. RecA1 and RecA3 were tagged with HA and βC1 with MBP. RecA1 and RecA3 protein size: ∼45 kDa, βC1: ∼59 kDa, MBP: ∼42 kDa.

To further verify the interaction of βC1 and RecA, we performed an *in vitro* pull-down assay (Figure S6A). The 6X-HIS-*Cv*RecA was transformed in BLR (DE3) cells and the lysate was passed through immobilized MBP or βC1 columns following elution and WB with anti-HIS. βC1, but not MBP, bound *Cv*RecA, suggesting that βC1 can selectively bind to RecA. Further, since RecA and βC1 both are ssDNA binding proteins, we analysed whether RecA is co-purifying with βC1 as a result of its tendency to bind to ssDNA. DNase I treated βC1 samples were also able to pull-down RecA, suggesting that the interaction of βC1 with RecA was not via DNA binding (Figure S6A). We further confirmed the interaction of βC1 with RecA by Affinity Purification Mass Spectrometry (AF-MS) (Table S1). RecA along with its other DDR counterparts such as RecBCD, dnaJ and dnaK were identified with high scores in AF-MS. Interestingly, nucleases like exonuclease-7 and SbcCD nuclease were also detected, reinforcing our previous observation of associated nuclease activity. These results suggest that RecA can physically interact with βC1 and is co-purified with βC1 during its recombinant expression in *E. coli*.

### RecA1, a plant homolog of bacterial RecA, can interact with βC1 *in planta*

It is not surprising that a bacterial protein such as RecA was able to interact with βC1, a geminiviral protein, since these viruses share similarities with bacterial systems and share evolutionary history (Koonin and Ilyina, 1992; Rekab et al., 1999; Krupovic et al., 2009; Kazlauskas et al., 2019). RecA has close homologs across all walks of life, we wondered if such an interaction is possible in host plants with functional consequences. RecA1 and RecA3 are the closest plant homologs of bacterial RecA. RecA1 is chloroplastic with a sequence similarity of 64%, whereas RecA3 is mitochondrial and 66% similar to *EcRecA* (Cerutti et al., 1992; Khazi et al., 2003; Rowan et al., 2010). It is important to note that although Rad51 type of proteins are considered as the functional homologs of bacterial RecA in eukaryotes, sequence-wise they are different from RecA (Figure S6B). Both RecA1 and RecA3 consist of a core ATPase domain with conserved Walker A and B motifs similar to RecA. Interestingly, plant homologs RecA1 and RecA3 are slightly larger than bacterial RecA with a distinct N-terminal region of ∼50 residues. Surprisingly, *Nt*RecA1 is larger and has only 77% sequence identity to *At*RecA1 (Figure 5B). We used a yeast two-hybrid system to identify if βC1 interacts with plant RecA homologs (Figure 5C and S6C). As expected, RecA exhibited strong interaction with βC1 in quadruple KO media. Previous study showed Rad51D interaction with Rep (C1) protein, hence we used Rad51D to check its interaction with βC1. Among the plant homologs *Nt*RecA1 exhibited strong interaction with βC1 while other homologs were unable to interact. To validate the Y2H results and to verify RecA1 interaction with βC1 *in planta*, we employed *in planta* pull-down assay (Figure 5D). Both *Nt*RecA1 and *Nt*RecA3 were tagged with HA and co-expressed with GFP-βC1. We detected interaction for *Nt*RecA1 in the pull-down assay but not *Nt*RecA3, suggesting that βC1 interacts specifically with *Nt*RecA1 (Figure 5D). Since complementing *Cv*RecA in BLR (DE3) expressing βC1 cells significantly reduced cell lethality, we tested whether the plant homolog of *EcRecA*, *NtRecA1*, which interacts with βC1 *in planta,* can complement BLR (DE3)::βC1 cells (Figure 5E). BLR (DE3)::βC1 cells, as expected, were not viable, but upon complementation with plant RecA1 showed a significant increase in cell viability. In agreement with pull-down assays, *Nt*RecA3 was able to complement βC1 only weakly (Figure 5E). All proteins expressed appropriately in the complemented system (Figure 5F). Together, these results suggested that βC1 can interact with RecA homologs in plants and such an interaction has conserved function since plant RecA1 was sufficient to alleviate βC1 induced genotoxicity in bacteria.

### DNA binding property of plant RecA1 is modulated by βC1

RecA1 is essential for maintaining the genetic stability of cpDNA. Binding of βC1 to *Nt*RecA1 might alter the activity of RecA1. To explore this further, we tested whether *Nt*RecA1 is capable of binding to DNA (Figure 6). *Nt*RecA1 was able to specifically bind to both ss and dsDNA probes (Figure 6A and B and S7A). Similarly, *Ec*RecA bound to both forms of DNA (Figure S7B). RecA is known to catalyze the branch invasion and migration at the site of DNA damage (Komori et al., 1999). We hypothesized that the DNA binding property of RecA might be altered in the presence of βC1. As SyYVCV βC1 has a strong affinity to ssDNA and rather a weak binding to dsDNA of smaller length, the assay was designed to probe the ability of RecA to bind dsDNA in the presence of WT βC1 or its truncation mutants. Interestingly, dsDNA binding of RecA was significantly altered with βC1 (Figure 6C). As shown earlier, the N-terminal half of βC1 was responsible for binding to RecA. Titrating (δC59) N-terminal half of βC1 protein with RecA reduced binding (Figure 6D), whereas, C-terminal half of βC1 (δN59) did not alter the DNA binding property of RecA (Figure S7C). These results suggest that βC1s interaction with NtRecA1 alters the dsDNA binding ability of the latter.

**Figure 6:**
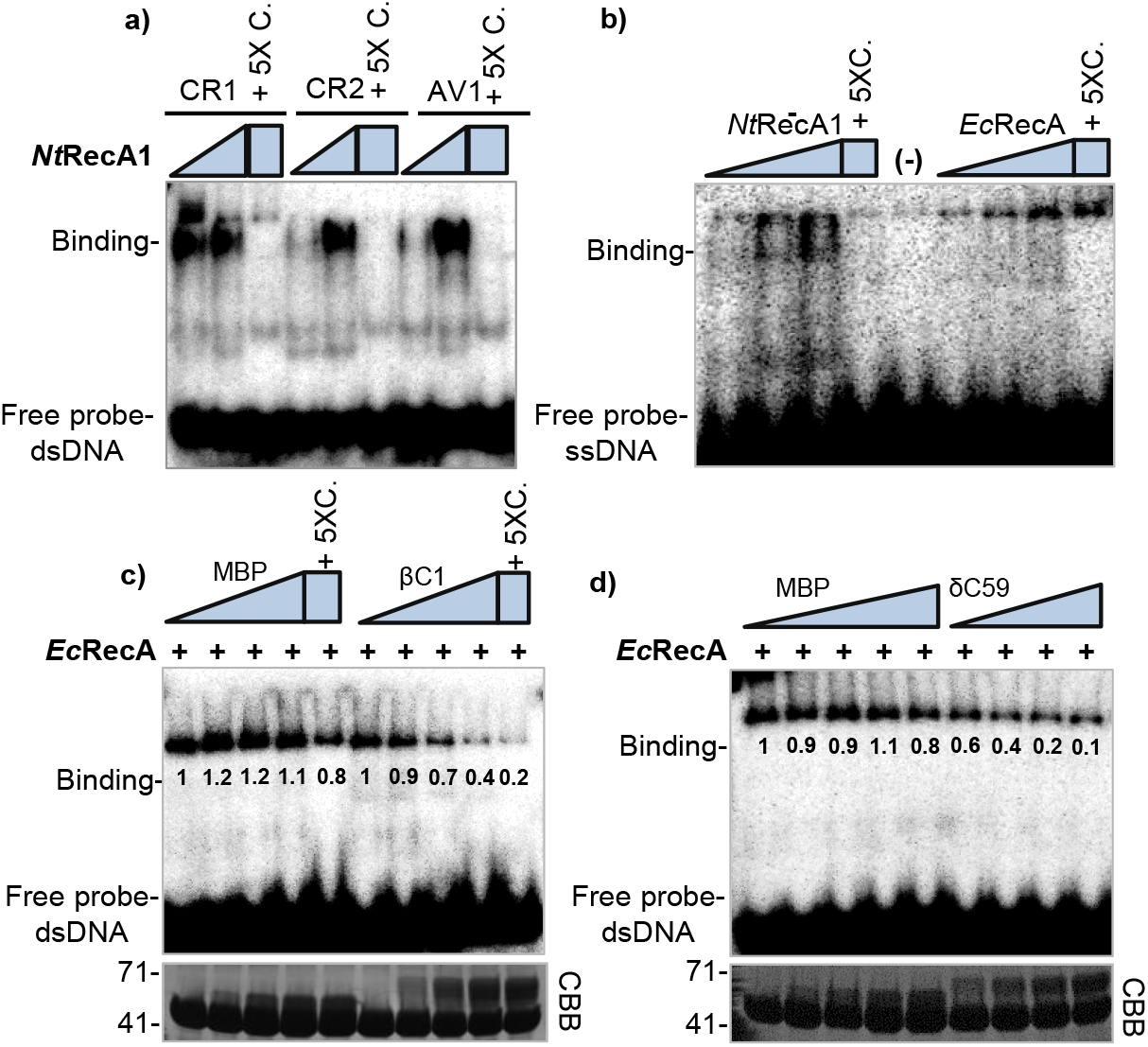
βC1 modulates DNA binding property of RecA. **a)** Gel shift assay showing interaction of *Nt*RecA1 with dsDNA probes. **b)** Same as a) except ssDNA probes. **c) and d)** EMSA competition assay: RecA protein was co-incubated with viral DNA probe along with varied concentrations of MBP-βC1, βC1 truncation mutant or MBP alone, before resolving. CBB shows amount of total protein. 5X represents concentration of competitive inhibitor (cold probe). MBP-βC1: 59 kDa, MBP: 42 kDa, RecA: 38 kDa.

### *Nt*RecA1 augments viral replication in host plants

*Geminivirus* replicative forms require assistance from host DNA repair machinery to form a complete genome (Singh et al., 2007; Yao et al., 2011; Richter et al., 2016). We tested whether the interaction of βC1 with *Nt*RecA1 alters viral replication by performing a viral replication assay (Figure 7A). p35S::*Nt*RecA1-HA and GFP-βC1 were co-infiltrated along with SyYVCV DNA-A in *N. benthamiana* and *N. tabacum*. Viral RFs were analysed using SB. As controls, p35S::GFP was used along with an empty vector. As expected and previously observed (Nair et al., 2020), βC1 enhanced viral titre (Figure 7A). In the presence of *Nt*RecA1 and WT βC1, but not with βC1-DM mutant, the accumulation of viral replicons increased. qPCR analysis of Rep (C1) region also suggested an additive effect of *Nt*RecA1 in the presence of βC1 but not with βC1-DM (C-terminal tagged βC1, functionally inactive) (Figure 7B). We also performed a similar experiment in *N. tabacum* where we used *Nt*RecA1 and *Nt*RecA3 in a time-course analysis of viral replication (Figure S8A). Viral SyYVCV DNA-A replication was higher in all the time-points in the presence of *Nt*RecA1 but not with *Nt*RecA3 (Figure S8A and B). These results suggested that the ability of βC1 to alter the DNA binding property of RecA1 is beneficial to viral replication.

**Figure 7:**
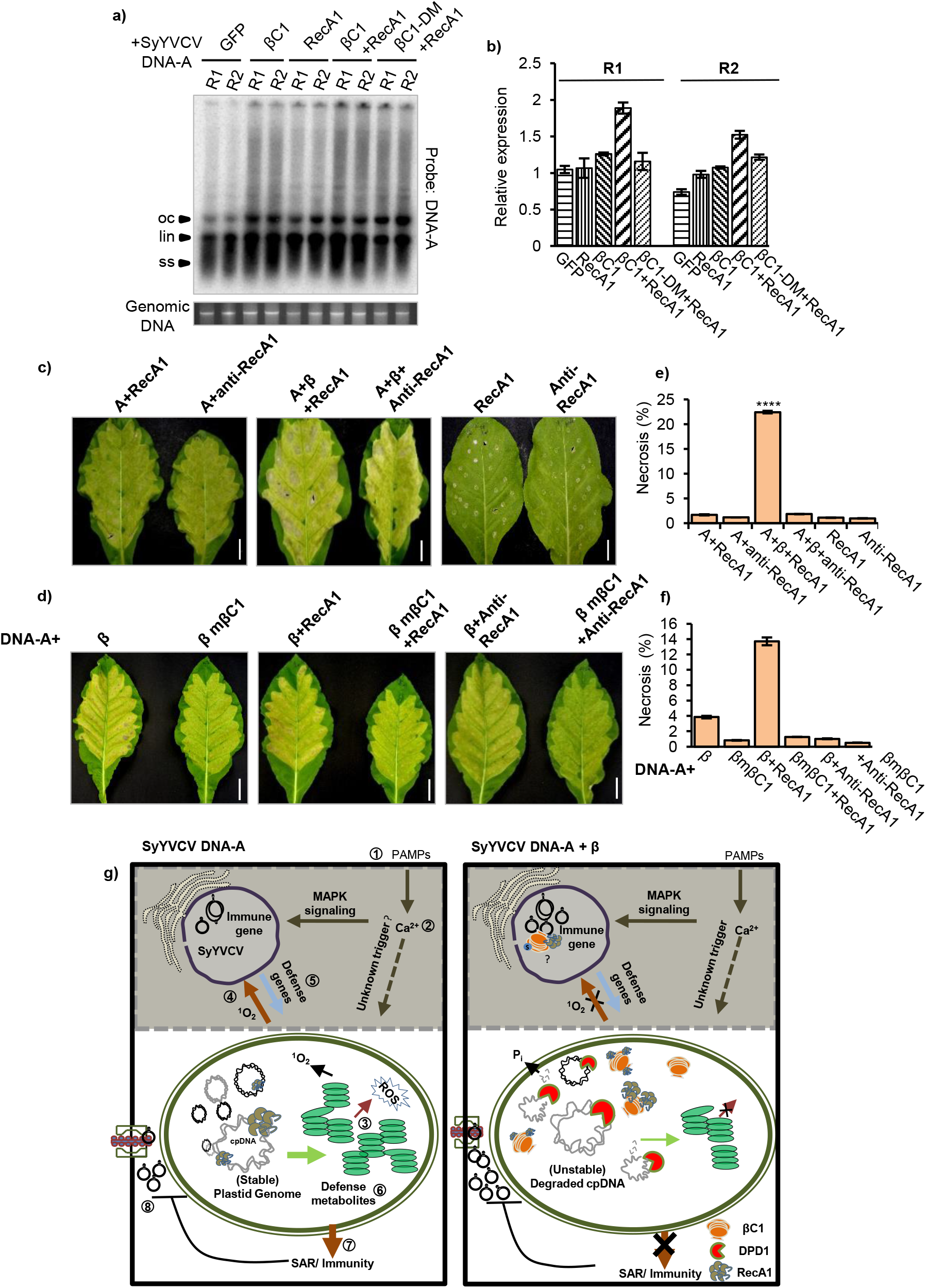
RecA1 augments pathogenicity determinant function of βC1. **a)** Viral replication assay with SyYVCV DNA-A partial dimer co-inoculated with *Nt*RecA1, βC1 or both, in *N. benthamiana*. SB blot was performed at 7DPI using full length DNA-A as probe. **b)** Same as a) except qPCR of viral Rep. *Actin* was used as internal control. R indicates biological replicates. **c)** RecA1 or anti-RecA1 infiltrated either alone or with DNA-A and DNA-A+β in *N. tabacum* leaves. N=3. **d)** Same as a) except DNA-βmβC1 was co-infiltrated along with RecA1 and anti-RecA1. d) and f) Quantification of the necrosis observed in infected leaves in a) and b), respectively. Quantification of necrotic area was done using FIJI. Scale bars in leaf is 2 cm. Images were taken at 12 DPI. **g) Summary: Left panel** (SyYVCV DNA-A) RecA1 maintains the multicopy plastid genome. 1) During virus infection, PRRs detects PAMPs. 2) Activated PRRs, phosphorylate special Ca^2+^channels leading to calcium influx. 3) The Ca^2+^ signal is relayed to chloroplast causing release of Ca^2+^ from thylakoids into the stroma. 4) Increased stromal Ca^2+^ leads to the increase in ^1^O_2_ which are the key molecules for retrograde signaling between nucleus and chloroplast. 5) ^1^O_2_ activates the transcription and translocation of nuclear defense responsive genes into the chloroplast. 6) Increased synthesis of hormones and secondary metabolite 7) These secondary metabolites travel through the plants inducing resistance and countering pathogenic invasion of new cells. 8) Intracellular viral movement is severely curtailed due to plastid mediated defense reducing symptom severity and disease. **Right panel:** In the presence of SyYVCV βC1, plastid genome maintainer RecA1 is recruited and forms a complex with βC1. Simultaneously, βC1 upregulates plastid nucleases like DPD1 to destabilize plastid genome, reducing the photosynthetic output and capability to code for key enzymes. In the presence of βC1, significantly reduced copy number of plastid genome severely hampers the retrograde signaling necessary for defense gene activation and immunity. Disruption of chloroplast defense signaling by βC1 cripples SAR and assists in viral infection.

### RecA1 enhances βC1-derived viral symptoms

βC1 is responsible for chloroplastic DNA degradation and chlorosis during infection (Figure 1). To analyse the function of its interaction with RecA1, we infected DNA-A or DNA-A+β in combination with PVX-RecA1 or PVX-antisense-RecA1 (anti-RecA1) (Figure 7C) on *N. tabacum* leaves. There was no difference observed in DNA-A-induced symptoms in the presence of either *Nt*RecA1 or anti-*Nt*RecA1 (Figure 7C, left panel). Interestingly, we observed an increase in chlorosis and large necrotic areas in DNA-A+β when co-infected with *Nt*RecA1 compared to anti-RecA1 (Figure 7C, middle panel and 7E). This result is in agreement with the results of viral replication assay. Neither *Nt*RecA1 nor anti-*Nt*RecA1 produced any chlorosis or necrosis when expressed alone (Figure 7C, right panel and Figure S8C), suggesting that *Nt*RecA1 can only augment symptom determinant function of βC1. Correspondingly, we observed an increase in some of the PVX-βC1 symptoms upon co-infection with RecA1 as compared to antisense-*Nt*RecA1 (Figure S8C and D). Additionally, large necrotic spots were observed in DNA-A+β but not in DNA-A+βmβC1 in the presence of *Nt*RecA1 (Figure 7D and F). Interestingly, DPD1 expression was upregulated in presence of WT βC1 irrespective of *Nt*RecA1 (Figure S8E). These results suggest that RecA1 directly or indirectly increases viral symptoms in the presence of DNA-β coding for βC1.

## Discussion

A mature plant cell contains hundreds of chloroplasts. Each chloroplast contains a relatively small-sized genome (cpDNA, 100-200 kb) in multiple copies (Day and Madesis, 2007; Sakamoto and Takami, 2018; Sato et al., 2003). In tobacco, Arabidopsis and in maize, cpDNA remains relatively stable until senescence (Golczyk et al., 2014). The high copy number of cpDNA is essential for adequate rRNA production required to sustain the arduous photosynthetic apparatus (Bendich, 1987; Udy et al., 2012). A significant decrease in cpDNA copy number severely affects the photosynthetic and metabolic process of the cell. Subunits of important photosynthetic enzymes are not abundant when compared to the cpDNA-coded mRNAs, suggesting the role of translational machinery as a checkpoint for chloroplastic efficiency. This observation suggests the importance of efficiently maintaining the cpDNA copy number during growth and development (Eberhard et al., 2002; Hosler et al., 1989). The relatively constant and high copy number of cpDNA is maintained by nucleoid-dedicated replication repair and organization proteins. A nucleoid proteome study identified a complex of 33 proteins involved in the homeostasis of cpDNA nucleoid (Majeran et al., 2012). The complex was enriched in replication (Polymerase IA, DNA Gyrase A and B etc.) and repair machinery (MutS, UV-REPAIR proteins (UvrB/C), PHOTOLYASES, and RecA orthologs). ssDNA and dsDNA breaks are very common in plant genomes due to the process of photosynthesis as well as their sessile nature (Noctor and Foyer, 2016; Bray and West, 2005; Fulcher and Sablowski, 2009). A dynamic and versatile DDR machinery to cope with such stresses is observed across plants (reviewed in (Manova and Gruszka, 2015). Homologs of bacterial RecA proteins are crucial for the maintenance of the organellar genome. RecA homologs, especially RecA1 forms complex with various DDR members to maintain the integrity of the plastid genome (Odahara et al., 2017; INOUYE et al., 2008; Odahara et al., 2015b, 2015a). Arabidopsis cp*recA* mutant lines had a drastic effect on the organelle function due to loss of genomic integrity and uncontrolled recombination in cpDNA. cpDNA copy number was further reduced upon removing other RecA orthologs like DRT proteins, suggesting an active role of RecA1 protein in the stability of the plastid genome (Rowan et al., 2010). It is not surprising that a pathogenicity determinant protein of a geminivirus, a group of viruses with archael origin, have evolutionarily conserved interactions with a DDR protein, such as RecA in bacteria and RecA1 in plants.

We observed pronounced misregulation of defense and hormone signalling pathways in transgenic βC1 expressing plants (Figure S1D). While these might have contributed to the phenotypes observed in these plants (Figure S1E), the differential expression of a set of DDR genes was surprising (Figure S2A). The deregulation of DDR genes in βC1 transgenic plants in the absence of viral infection indicated the direct role of βC1, independent of viral replication. Interaction of βC1 with viral genome might direct DDR proteins to viral DNA and or, βC1 might play a direct role in the genomic instability of host cells. Interestingly, in our transcriptome data, we also observed a large number of chloroplastic genes misexpressed. SyYVCV βC1 localizes to the chloroplast and chlorosis was observed only in the presence of βC1. As expected, plastid DNA levels reduced in these plants and indicated a clear role for βC1 in organellar genome instability (Figure 1).

The ability of βC1 to induce genomic instability was due to its DNA binding ability and an associated nuclease activity, the latter due to its interacting nuclease partners. βC1 associated nuclease activity led to induction of genotoxic stress in bacterial and plant cells deficient in DDR functions (Figure 2). We assume, in repair proficient cells, βC1 induced genotoxic stress is countered by cellular DDR machinery, allowing the affected cells to survive. In agreement with this, βC1 expressing cells did not survive a sub-lethal UV or bleomycin treatment. The genotoxic activity of βC1 in *E. coli* can be extrapolated to its role during plant infection, where plastid DNA degradation and cell death are associated with its expression. Additional sub-lethal stress was essential for tipping the balance of repair and damage in genotoxicity assay in *E.coli* cells, whereas in plants, replicating virus might act as the additional stress. Based on our observations, βC1 directly appears to inhibit RecA1 from protecting cpDNA and possibly redirects RecA1 for virus replication. Simultaneously, βC1 induced a plastid specific nuclease, DPD1, to induce cpDNA degradation. DPD1 is a Mg^2+^ dependent exonuclease well known to degrade organellar DNA (Sakamoto and Takami, 2018).

Reduction in plastid genome results in decreased coding capacity and perturbation in the structure of chloroplast, significantly altering its antiviral response-ability. As a consequence, SA induced PATHOGENESIS RELATED GENE (PR) expression is severely repressed. SA is synthesized in chloroplast and is regulated by plastid retrograde signalling mediated by ROS. βC1 selectively disrupts plastid signalling, down-regulating ROS as was evident in βC1 transgenic plants. The mβC1 without genotoxic activity neither affected plastid DNA nor was able to repress PR gene expression (Figure S8F and G). Alternatively, as geminivirus infects differentiated cells, availability of raw materials for replication is limited. Degradation of the plastid genome by DPD1 might be a common mechanism to cope with phosphate deprivation (Takami et al., 2018) and might be the key player of active salvage pathway (Tang and Sakamoto, 2011). Nucleic acids and phospholipids are the most abundant source of phosphorous esters. Multiple copies of the organellar genome can encrypt massive storage of phosphate that a virus might exploit during replication. In accordance with the hypothesis, along with DPD1, we also observed significant upregulation of bifunctional nuclease (BFN1) that are well characterized for their role in salvaging RNA during senescence (Figure S2A) (Pérez-Amador et al., 2000). Organellar DNA serves as a “dispensable” phosphate source that can be readily tapped into without significantly disturbing the cellular homeostasis. Nutrient stress leads to reduction in organellar DNA and relocation of vital minerals, similar scenario should also exist in cell-cycle exited cell during viral replication. Such possibilities are interesting and might be of future interest in other plant-pathogen interactions. It is worth mentioning that most plant pathogens induce chlorosis and necrosis in susceptible host plants.

The interaction of βC1 with RecA1 appears to have a profound effect on latter’s function (Figure 6). The impact of this modulation of the dsDNA binding ability of RecA bound with βC1 is currently unknown. Since this interaction of βC1 with RecA1 has functional significance in terms of augmenting viral titre and infection symptoms, for example, chlorosis and necrotic symptoms were enhanced in the presence of βC1 and RecA1, it is clear that this interaction helps βC1 as a pathogenicity determinant. In agreement with this, upon co-infection of βC1 with RecA1, we observed an increase in βC1-induced symptoms. However, the exact functional significance of the interaction between βC1 and RecA1 is yet to be understood.

The cellular repair pathway plays an important role in maintaining the integrity of the nuclear as well as organellar genome (Kunkel, 2004). We observed recombination proteins like RecA-cluster (RecBCD, RecQ, and RecF), dnak and dnaJ in βC1 AF-MS, suggesting interaction and modulation of the DDR machinery by βC1 (Table S1). In plants, *Nt*dnaJ an important DDR member, was transcriptionally upregulated in βC1 transgenic lines (Yamamoto et al., 2005; Majeran et al., 2012). Considering these evidences, it is also highly likely that βC1 is directly involved in replication and repair of virus RFs, guiding cellular machinery to viral replisome in host nucleus.

Here we identified a DDR protein RecA1 as a functional partner of viral pathogenicity protein βC1. Other than its observed functions in the chloroplast that we elucidated here, βC1 might also recruit RecA1 into the viral replication complex to assist in the RF repair process. Evidence for this aspect of the interaction between RecA1 and βC1 needs further experimentation.

## Methods

### Plasmids and cloning

Cloning was performed as previously described (Nair et al., 2020). Briefly, the partial dimer of DNA-A and DNA-β was amplified from pSD30 and pSD35, respectively, and cloned into pBIN19 using BamHI and SacI sites (Das et al., 2018). For plant transformation and transient expression, βC1 and RecA proteins were cloned into modified pBIN19 (35S CAMV promoter) vectors using BamHI and SacI sites for βC1, and BamHI and XhoI for RecA proteins. For βC1 fusion constructs, βC1 was amplified using primers AN34 and AN35 containing SalI and SacI (Table S2). eGFP was amplified from pMEL2 (pBIN HSP30-eGFP) using primer pair AN30 and AN31 having BamHI and SalI along with a short linker sequence (Gly-Gly-Ser-Gly). The vector pBIN-GFP was digested with BamHI and SacI to release ∼700-bp product. βC1 amplicon was digested with SalI and SacI and eGFP amplicon was digested with BamHI and SalI. A three fragment ligation was performed using βC1 fragment, eGFP fragment, and linearized vector. For recombinant protein purification vectors, the pMAL-p5E vector has an AMPr resistance gene and a maltose-binding protein (MBP) ORF driven by a TAC promoter. The MCS is at the C-terminus of MBP protein just after an enterokinase cleavage site. βC1 (insert) was amplified from pAN3 using primer pair AN3 and AN4 containing KpnI, precision protease cleavage site (Leu Glu Val Leu Phe Gln/Gly Pro) in AN3 and XhoI, NotI sites in AN4. pMAL-p5E vector and βC1 were digested with KpnI and NotI. The substitution mutations in βC1 were generated using overlapping PCR primers harbouring the required modification or by site-directed mutagenesis kit (Invitrogen). Primer and plasmid used in the study are detailed in (Tables S2, S3 and S4).

### Recombinant protein purification

For protein purification, Bl21 (DE3) or Rosetta-Gammi (DE3) (Novagen) cells were typically used unless otherwise mentioned. For purification of βC1 and its mutants, cells were grown to OD 0.7 at 37°C and induced with 0.3 mM IPTG at 20°C. The induced culture was incubated with shaking at 16°C for 18 h. The cells were pelleted and lysed using sonication (10 sec on and off for 15 cycles, 60% amplitude) in the lysis buffer (50 mM Tris-Cl pH-8, 500 mM NaCl, 5% glycerol, 5 mM 2-mercaptoethanol, Igepal 0.01%, and protease inhibitor tablets (Roche)). The lysate was clarified using centrifugation and the supernatant was passed through pre-equilibrated dextrin-sepharose beads (GE). Non-specific protein binding was washed off using high salt washes and the bound protein was eluted with 12 mM maltose. The eluted protein was concentrated using amicon filters (Millipore). The concentrated protein was then passed through a Q-sepharose (Ion-exchange column, GE) and protein was eluted using NaCl gradient. The ion-exchanged purified protein was further concentrated and passed through a size-exclusion column (SD-200, HiLoad 16/600 200 pg superdex preparative column (GE)) and the protein fraction was concentrated and stored in storage buffer (25 mM Tris-Cl pH-8, 100 mM NaCl, 5% glycerol) at −80°C.

### Transgenic plants and transient expression

Transformation of tobacco (*N. tabacum*, Wisconsin 35) was performed as described previously (Sunilkumar et al., 1999; Nair et al., 2020). Briefly, leaf discs prepared from 3 week-old *N. tabacum* plants were infected with *Agrobacterium* strain LBA4404 (pSB1) harboring genes of interest (Stachel and Zambryski, 1986; Yanofsky et al., 1986). Transformants were selected on a kanamycin medium. Transient over-expression was performed on 3 to 4 week old *N. tabacum* leaves using *Agrobacterium* LBA4404 (pSB1) strains having appropriate genes, suspended in an infiltration buffer (10 mM MES, 10 mM MgCl_2_, pH 5.7, and 100 μM acetosyringone). A culture of 0.6 OD was used for infiltration unless otherwise mentioned. The resuspended culture was incubated for 1 h before infiltration. For protein expression studies and replication assays a 1:1 ratio of the culture of the same OD was premixed and then infiltrated using a needle-less 1 ml syringe onto 2nd whorl of leaves from the top of either *N. tabacum* or *N. benthamiana*. Samples were collected only from the infiltrated area of the leaves in all samples.

### RNA sequencing analysis

RNA sequencing analysis was done as previously described (Jha et al., 2021). Paired-end (100 × 2) RNA-seq reads were adapter trimmed using CUTADAPT (Martin, 2011) and aligned to the genome (*N. tabacum*: TN-90) using HISAT2 (Kim et al., 2019). Differentially expressed genes were identified using Cuffdiff with log2 fold change > 1.5 (Trapnell et al., 2011), and GO analysis was performed using Panther (Mi et al., 2021).

### Plant growth conditions

For phenotyping of plants, 2-week old rooted plants grown in rooting media were transferred to soil and kept in a controlled environment (growth-chamber, temperature: 24°C, light: 4 LSI light with 12 h cycle and RH 70%) for 1 week to acclimatize. Hardened plants were further transferred to larger pots in transgenic green-house (temperature: 24°C, RH 70-80% and natural light cycle).

### Plant total protein isolation and western blotting

Total protein was isolated using the acetone-phenol extraction method (Wang et al., 2006; Nair et al., 2020). Briefly, 200 mg of tissue was finely ground in liquid nitrogen, and protein was precipitated by 10% TCA (trichloroacetic acid, Sigma) in acetone, The resultant pellet was washed with 0.1 M ammonium acetate in 80% methanol followed by 80% acetone wash. The acetone extracted pellet was briefly dried to remove excess acetone and further extracted with 1:1 ratio SDS extraction buffer (850 mM sucrose, 100 mM Tris-Cl pH-8, 750 mM 2-mercaptoethanol, and 2% SDS) and phenol (pH-8.0 Tris-saturated). The supernatant was precipitated overnight using 0.1 M ammonium acetate in 100% methanol at −20°C. Precipitated protein was pelleted (13000 rpm for 30 min) and was washed once with 100% methanol and then with 80% acetone, air-dried, and resuspended in 2X SDS lamelli sample loading buffer containing 6 M urea and 1% CHAPS.

For western blot analysis, 20 µg of total protein was loaded either on a 12% Tris-Glycine SDS gel or 4-20% Bio-Rad precast gels. The gel was resolved under constant voltage (100 V). Resolved proteins were transferred to a nylon membrane (GE (Amersham) Protran, 0.2 μm) using the Bio-Rad protran transblot apparatus. The transfer was performed for 120 min at a constant voltage of 90 under the ice with transfer buffer maintained at 18-20°C. The blot was washed with TBST (TBS with 0.1% tween 20) and was blocked with either 5% blotting grade blocker (Bio-Rad) or 4% BSA (Sigma) in TBST. Blocked membranes were probed for the protein of interest using specific antibodies and imaged using Image quant 4000 LAS in chemiluminescence mode (GE). Stripping of blots was done at room temperature (Restore western stripping buffer, Thermo Fisher) according to the manufacturer’s instructions. Bands were quantified and normalized using FIJI software. Antibodies used in this study are listed in Table S5.

### Immuno-precipitation

Immuno-precipitation was performed as previously described (Nair et al., 2020). Briefly, Infiltrated or transgenic plant leaves expressing the protein of interest were finely powdered under liquid nitrogen. Three volumes of lysis buffer (50 mM Tris-Cl pH 7.4, 150 mM KCl, 1% Triton-X100, Protease inhibitor 1 X [Roche], NEM 20 μM) was added to 2 g of powdered tissue. The lysate was clarified by high-speed centrifugation and incubated with GFP-Trap (Chromtek) for 3h at 4°C. Beads were magnetically separated from the lysate and washed 5 times in wash buffer (50 mM Tris-Cl, pH 7.4; 150 mM KCl, 1 mM PMSF) until the green colour completely disappeared. The washed beads were transferred to a 1.5 ml tube and again washed twice with wash buffer. The buffer was completely removed and 3X SDS sample dye was added to the beads and incubated at 70°C for 10 min. The pull-down products were resolved in 4-20% Tris-Glycine SDS gradient gels (Bio-Rad).

### Mass-Spectrometry

In-solution digestion of the purified protein was performed using 13 ng µl^−1^ trypsin in 10 mM ammonium bicarbonate containing 10% (v/v) acetonitrile. Approximately 50 ng of the prepared samples was subjected to liquid chromatography/tandem mass spectrometry (LC-MS/MS), using a LTQ Orbitrap XL (Thermo Scientific), HCD activation, C-18 column, 15 cm length. The data was analysed using Proteome Discoverer (Thermo Scientific). For peptide identification, Sequest HT search engine was used against the combined target-decoy database with the following parameters: enzyme: trypsin; maximum missed cleavage: two; variable modifications: oxidation. Search tolerance parameters were as follows: minimum peptide length; 6, maximum; 144, false discovery rate, <1%.

### Viral replication assay and Southern blotting

Viral titre assay was performed as previously shown (Shivaprasad et al., 2008, 2006; Nair et al., 2020). Partial dimer of SyYVCV DNA-A, DNA-β, and 35S driven plasmids were mobilized into *Agrobacterium* strain LBA4404 (pSB1) and co-infiltrated alone or in various combinations into *N. tabacum* leaves. Genomic DNA from infiltrated and systemic leaves were isolated using the CTAB method (Rogers and Bendich, 1994). An equal amount of genomic DNA normalized using Qubit and gel-based quantification was loaded onto a 0.7% TNE agarose gel and resolved at 5 V/cm. The transfer was performed as previously mentioned (Shivaprasad et al., 2006) and blots were probed with full-length DNA-A in case of replication assay and psbM gene probe (3-kb) for plastid Southern blot. The probes were internally labelled with dCTP alpha P32 (BRIT, India) using the Rediprime II kit (GE). Blots were scanned using Typhoon Trio Scanner (GE) in phosphorescence mode.

### Yeast two-hybrid transformation and screening

Yeast transformation was performed as described with minor modifications (Gietz and Woods, 2002). Freshly streaked AH109 cells were used to initiate primary culture grown overnight in YPD media (yeast extract 1%, bacterial peptone 2%, and dextrose 2%). Cells were grown to A600 = 0.6 OD. About 10 ml of cells were pelleted per transformation. The freshly pelleted cells were transferred to a 1.5 ml centrifuge tube and washed with deionized sterile water followed by 0.1 M lithium acetate. Transformation mixture (PEG 3000 50%, salmon sperm DNA and lithium acetate) was added to the washed cells, and cells were resuspended. The corresponding mixture of AD and BD plasmids was added to the transformation mixture followed by vortexing for 30 s. The mixture was incubated for 30 min at 30°C and 30 min at 42°C. The reaction mixture was removed and cells were resuspended in 2 ml YPD media and allowed to recover for 2h before plating onto an auxotrophic media. All Rec and Rad proteins were translationally fused with the Activation Domain (AD) of pGADT7 AD (Takara Bio). βC1 was fused with Binding Domain (BD) and cloned into pGBKT7 BD (Takara Bio). Plasmids were transformed into AH109 strain as described previously (Gietz and Woods, 2002). Transformants were screened on -Leu, -Trp media followed by screening for interaction on -Leu,-Trp, -His with or without 3-AT (Sigma-Aldrich).

### Electrophoretic Mobility Shift Assay (EMSA)

EMSA was performed as previously described (Csorba and Burgyán, 2011). Briefly, oligos were end-labelled using T4 polynucleotide kinase (NEB) with γ-32P. Labelled oligos were diluted as mentioned for each experiment typically to 100 to 200 pg. Labelled oligos were incubated with protein in EMSA binding buffer (50 mM Tris-Cl pH-8, 100 mM NaCl, 5% glycerol) for a specific time interval as detailed in the experiment followed by stopping the reaction by addition of non-denaturing stop-dye. The reaction was further resolved in an 8% native TBE gel and exposed to a phosphor screen for development. The phosphor screen was scanned using Typhoon trio plus (GE) and the image was analysed using FIJI. For nuclease assay, buffer contained 50 mM Tris-Cl pH-8, 100 mM NaCl, 5% glycerol and 5 mM Mg^2^

## ACKNOWLEDGMENTS

We thank members of Shivaprasad lab for suggestions. We thank the Next Generation Genomics, radiation, CIFF and mass-spec at NCBS-TIFR, Bangalore. We thank Dr. Anjana Badrinarayanan and Afroze C. for critical discussion on RecA function and for cvRecA antibody. We also thank Prof. K. Veluthambi for binary vectors and *Agrobacterium* strains. This study was supported by Department of Atomic Energy, Government of India, under Project Identification No. RTI 4006 (1303/3/2019/R&D-II/DAE/4749 dated 16.7.2020). ANN acknowledges a fellowship from DBT, India. This work was also supported by NCBS-TIFR core funding and grants (BT/PR12394/AGIII/103/891/2014;BT/IN/Swiss/47/JGK/2018-19; BT/PR25767/GET/119/151/ 2017) from Department of Biotechnology, Government of India.

## AVAILAIBLITY OF DATA AND MATERIALS

All data generated or analysed during this study are included in this published article (and its supplementary information files) Source data for all the images, gels and blots used in the study are provided as original raw file, and images or blots used in the figure files are also provided as uncropped images with relevant area labelled. RNA-Seq data is available under GEO accession number: GSE189526.

## AUTHOR CONTRIBUTIONS

AN and PVS designed the study, analyzed the data and wrote the manuscript. AN performed almost all the experiments. HCY helped with PVX and tobacco transgenics. ANN helped with transcriptome analysis.

---

**Figure S1:**
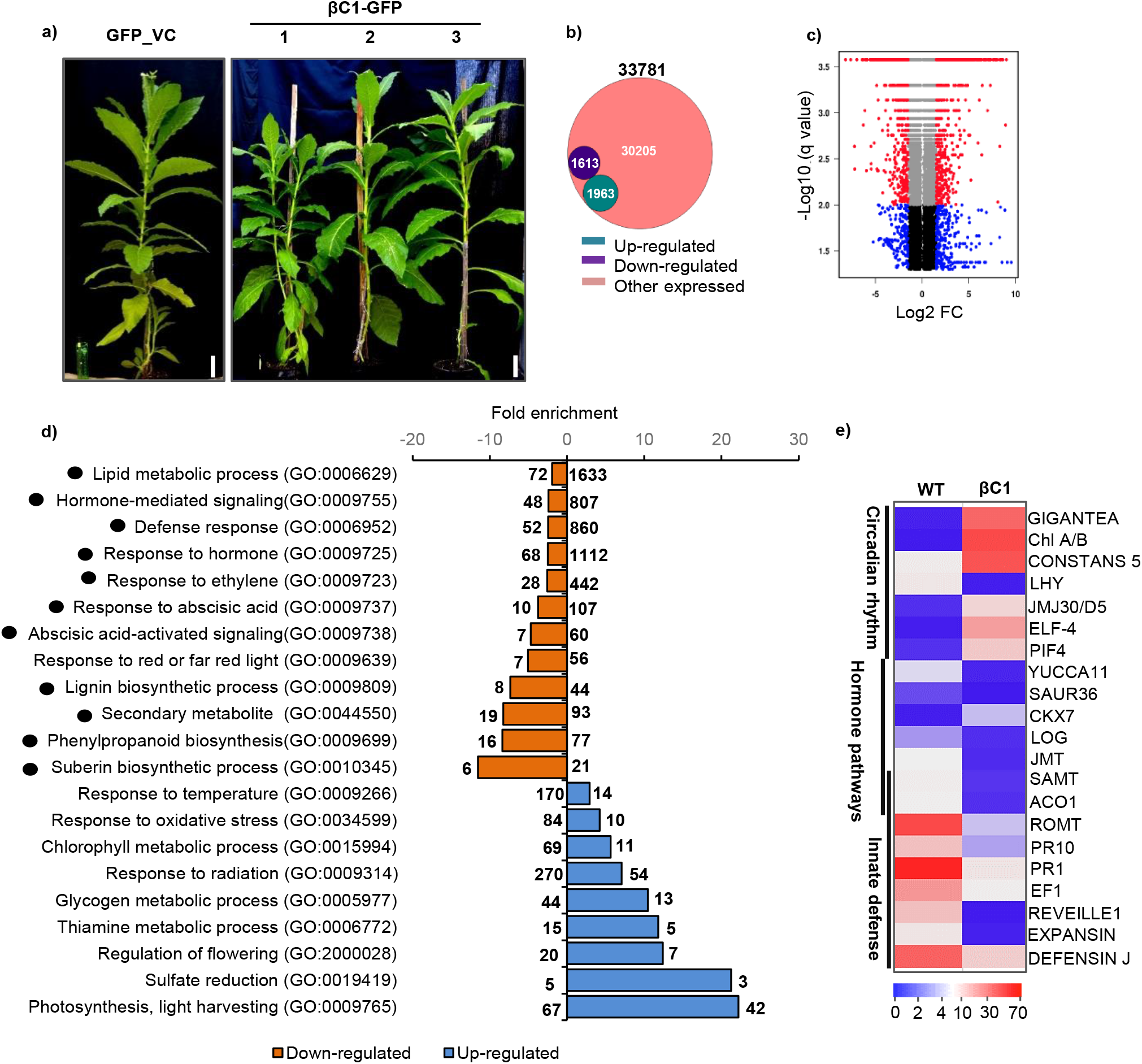
Signalling pathways are deregulated in βC1 transgenic lines. **a)** *N. tabacum* transgenic plants over-expressing C-terminal GFP tagged βC1 showing similar lack of phenotype to vector plants as reported in previous studies (Cheng et al., 2011). n=2 for each transgenic line. b) Venn schematic showing the number of DE loci in βC1-OE plants. **c)** Volcano plot showing DE loci in βC1-OE plants. **d)** DE loci of βC1 transgenic line (βC1_6) characterized under representative GO (Gene Ontology) classification. Numbers inside the bar graph represents enriched DE loci and numbers on top of respective bars indicate total number of reference loci in each process. Black circle represents defense pathway processes. **e)** Heat map showing normalised FPKM values of major pathway genes in βC1 plant versus vector control plants. Size bar in a) is 5.8 cm.

**Figure S2:**
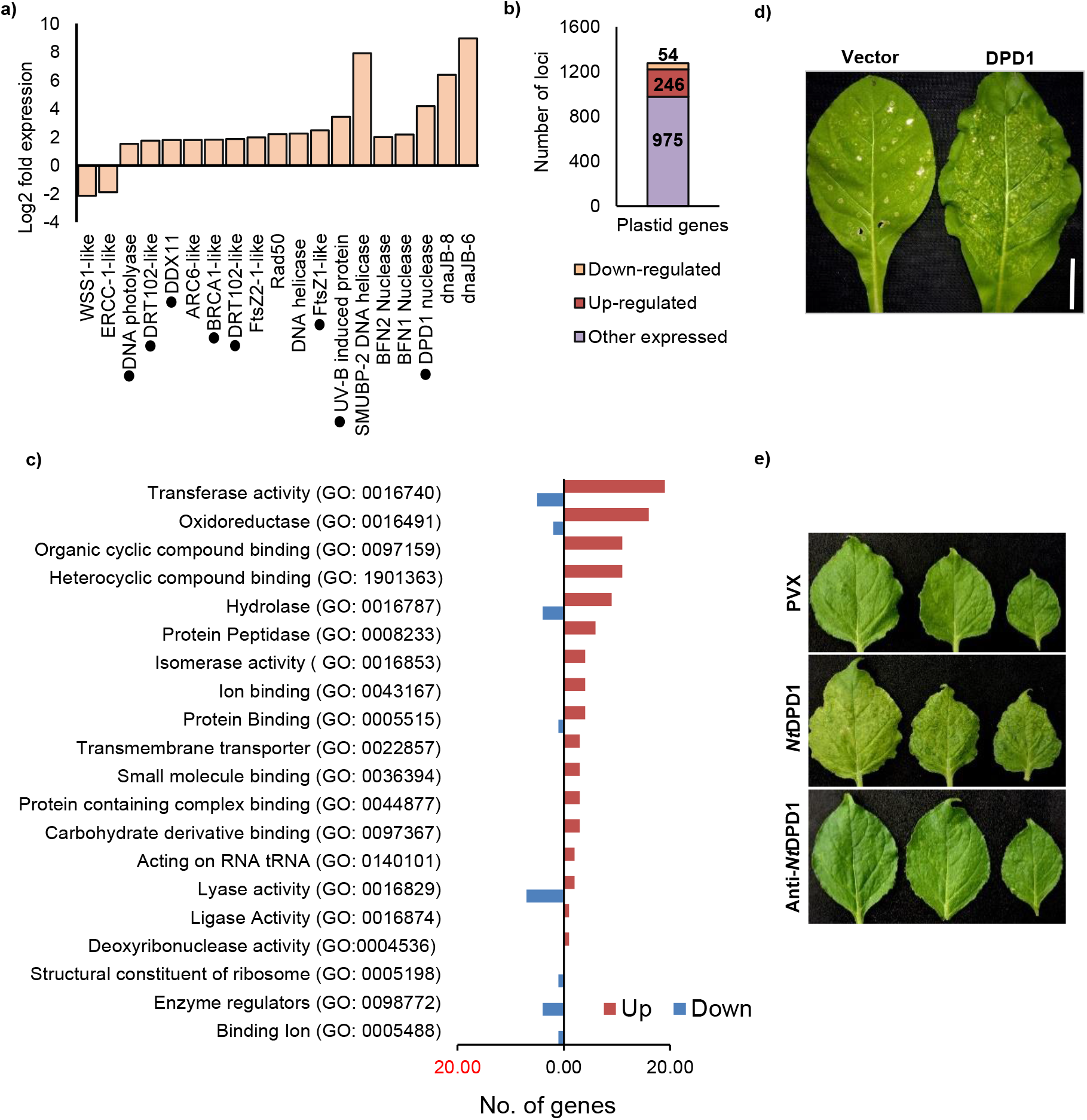
Plastid localised genes are significantly misexpressed in βC1-OE lines. **a)** DE of important DDR pathway genes in βC1 transgenic lines. **b)** Plot showing DE loci coded by chloroplast genome in βC1-OE plants. **c)** DE loci characterized under representative GO (Gene Ontology) processes for chloroplastic genome of βC1-OE plants. **c)**. Black circle represents known chloroplastic localization status. **d)** Phenotype of the *N. tabacum* leaf infected with PVX-*NtDPD1* or PVX. 8 DPI, local leaf, n=4 plants. **e)** Same as d) except *N. benthamiana* plants were infected with PVX-*NtDPD1*, PVX-Anti-*Nt*DPD1 and PVX. 8 DPI, systemic leaves, n=5 plants. Black circle represents known chloroplastic localization.

**Figure S3:**
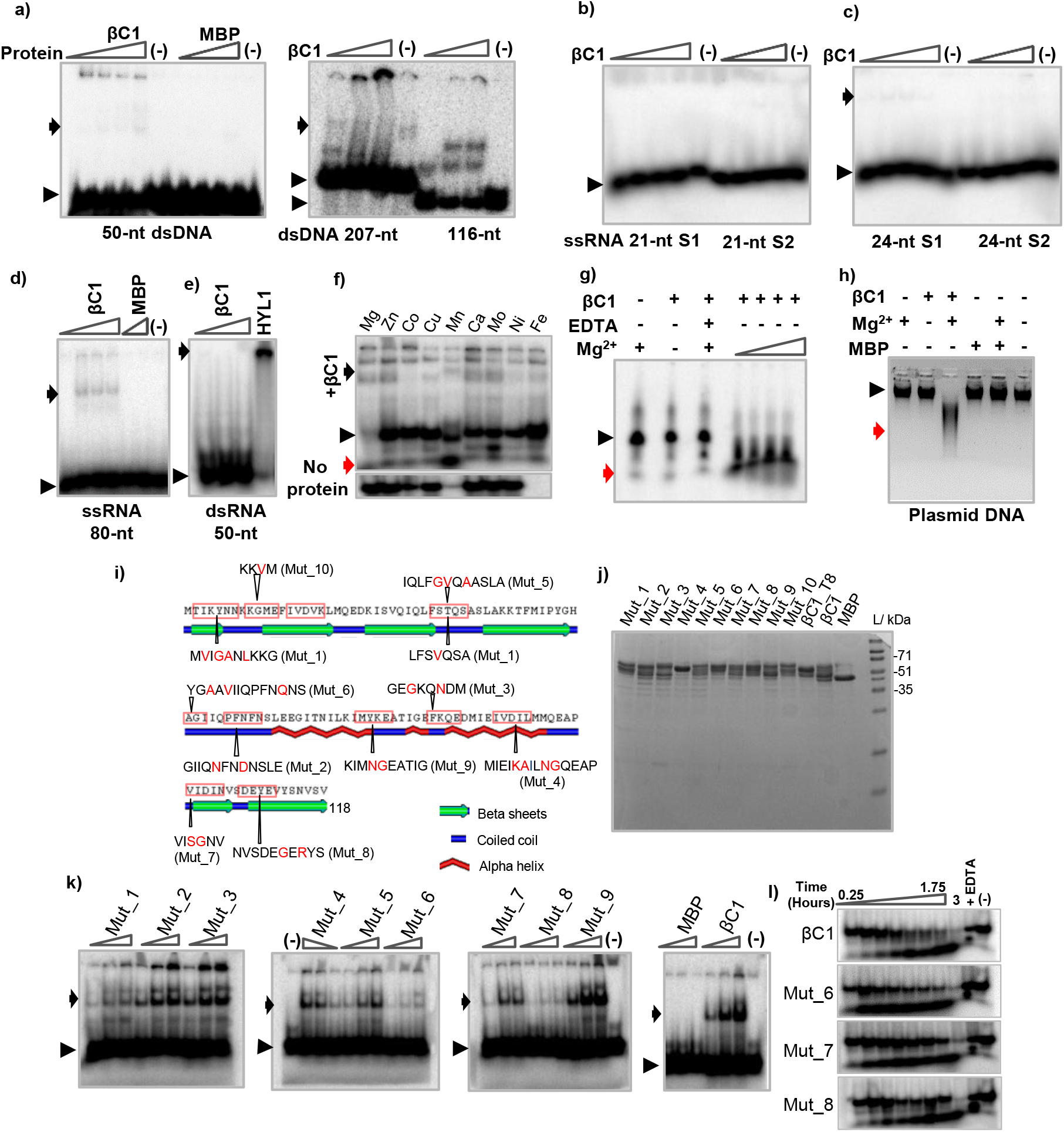
βC1 binds preferably to ssDNA *in vitro*. **a)** Binding of βC1 to dsDNA. **b)** EMSA of βC1 with ssRNA (21-nt). S1 and S2 are two different probes. **c)** same as b) except using 24-nt ssRNA. **d)** EMSA of βC1 with long ssRNA (80-nt). **e)** EMSA of βC1 to dsRNA (50-nt). **f)** Co-factor dependency. Various cations (1 mM) were incubated with βC1 to analyze the effect on ssDNA degradation. Mn^2+^ and Fe^2+^ precipitated the DNA in no protein control. **g)** Mg^2+^ dependency of ssDNA nuclease activity associated with purified MBP-βC1, denaturing PAGE. **h)** Nuclease assay with 12-Kb plasmid DNA. **i)** Schematic representation of SyYVCV βC1 showing scheme for point mutations. Residues indicated in red are substituted in mutants **j)** CBB gel showing recombinantly purified βC1 mutants depicted in i). **k)** *in vitro* binding assay: gel showing binding affinity of βC1 mutants towards 49-nt ssDNA (with EDTA, no Mg^2+^). **l)** Time point nuclease assay: mutant 6, 7 and 8 was incubated with ssDNA in nuclease buffer containing Mg^2+^. All the binding experiments were repeated multiple times with βC1 and mutants proteins extracted in biological replicates. A representative image of the binding experiment is shown here. (-) indicates no protein. Other details are same as figure 2.

**Figure S4:**
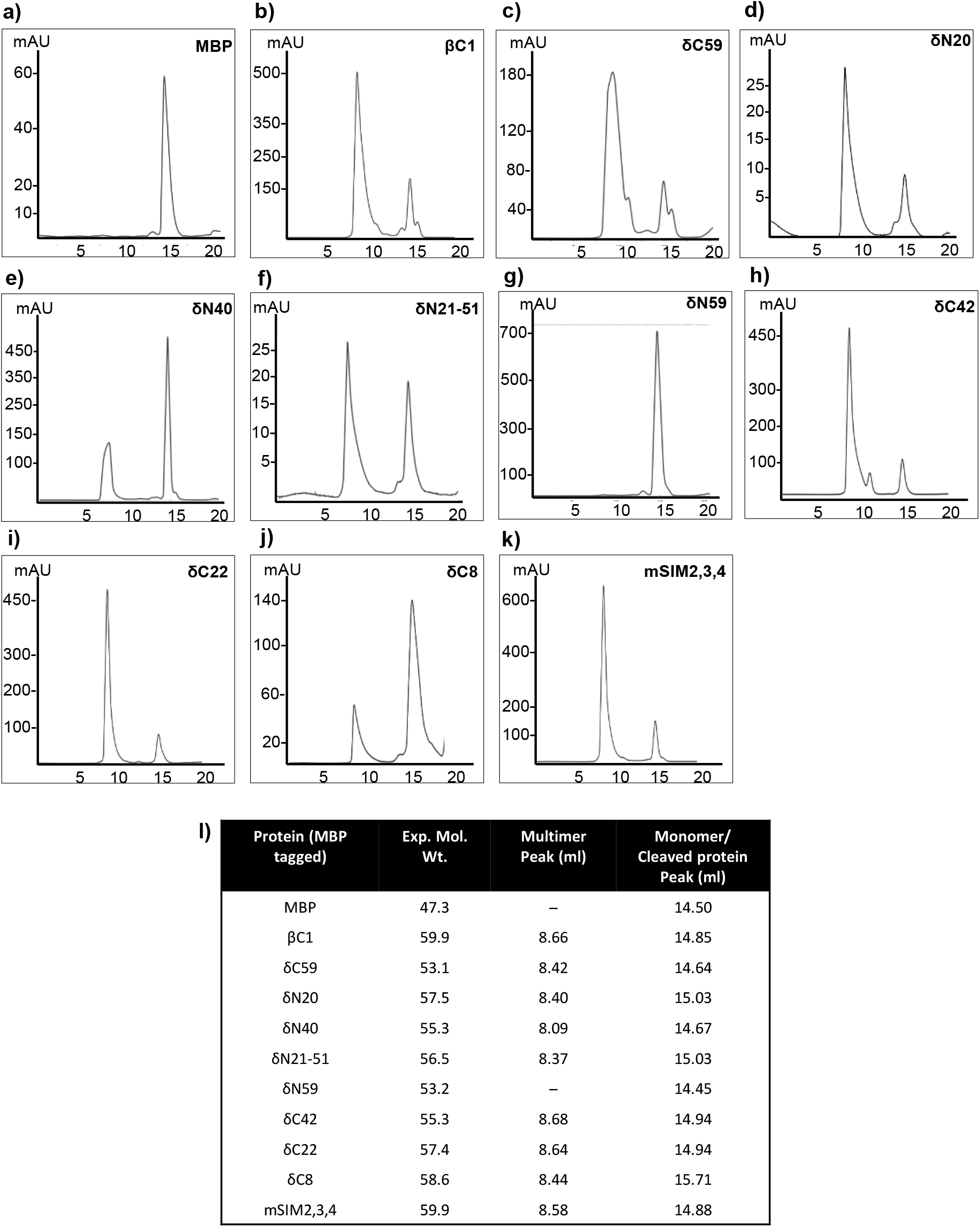
Size exclusion profile of βC1 and its truncation mutants. **a) to k)** SD-200 Superdex 10/300 GL analytical columns were used for the assay. All proteins were MBP tagged. a) MBP, b) βC1, c) δC59, d) δN20, e) δN40, f) δN21-51, g) δN59, h) δC42, i) δC22, j) δC8, k) mβC1 (mSIM2,3,4). **l)** Table indicating elution volume of each peak for individual proteins. All the assays were biologically replicated multiple times in SD200 as well as SD75 Superdex column. N=3, biological replicates (separate protein batches) on SD-75 columns.

**Figure S5:**
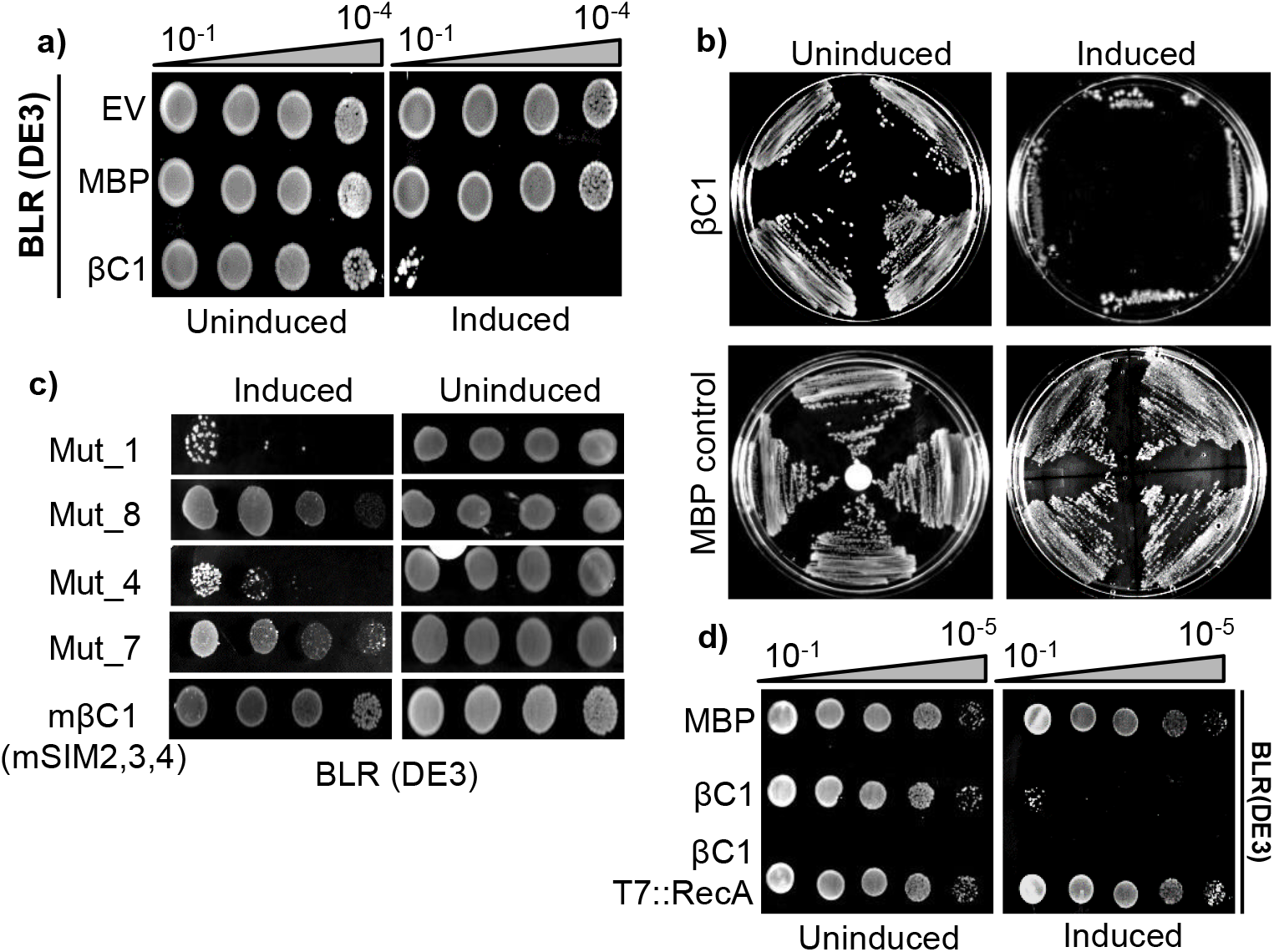
Plant *RecA, a Rad51* homolog, is essential for cell survival in presence of βC1. a) DNA damage sensitivity assay: βC1 was expressed in BLR (DE3) cells (*RecA* deficient) followed by induction with 0.1 mM IPTG. **b)** Same as a) but streak assay. **c)** Growth assay of BLR (DE3) cells expressing different mutants of βC1. **d)** Same as b), but with *Caulobacter RecA* (*Cv*RecA) complemented in BLR (DE3) cells co-expressing βC1. EV indicates empty vector. All the assays were biologically replicated at least 3 times.

**Figure S6:**
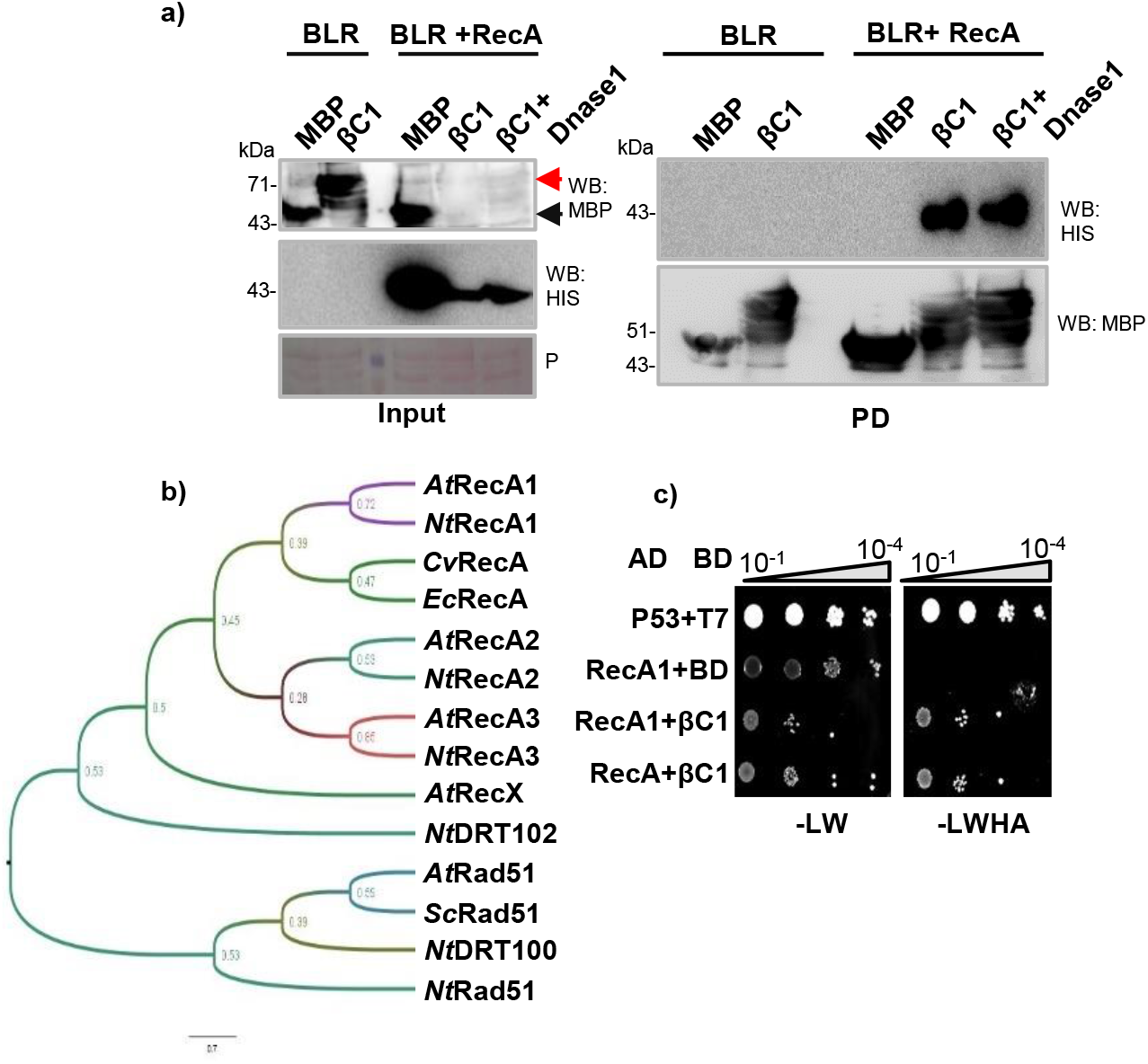
RecA directly interacts with SyYVCV βC1. **a)** *in vitro* pull-down assay with 6X-HIS-RecA and MBP-βC1. **b)** Maximum likelihood (ML) tree of RecA homologs based on core ATPase region. Numbers indicate bootstrap values on nodes. **c)** Y2H assay showing interaction of βC1 with RecA and its plant homolog.

**Figure S7:**
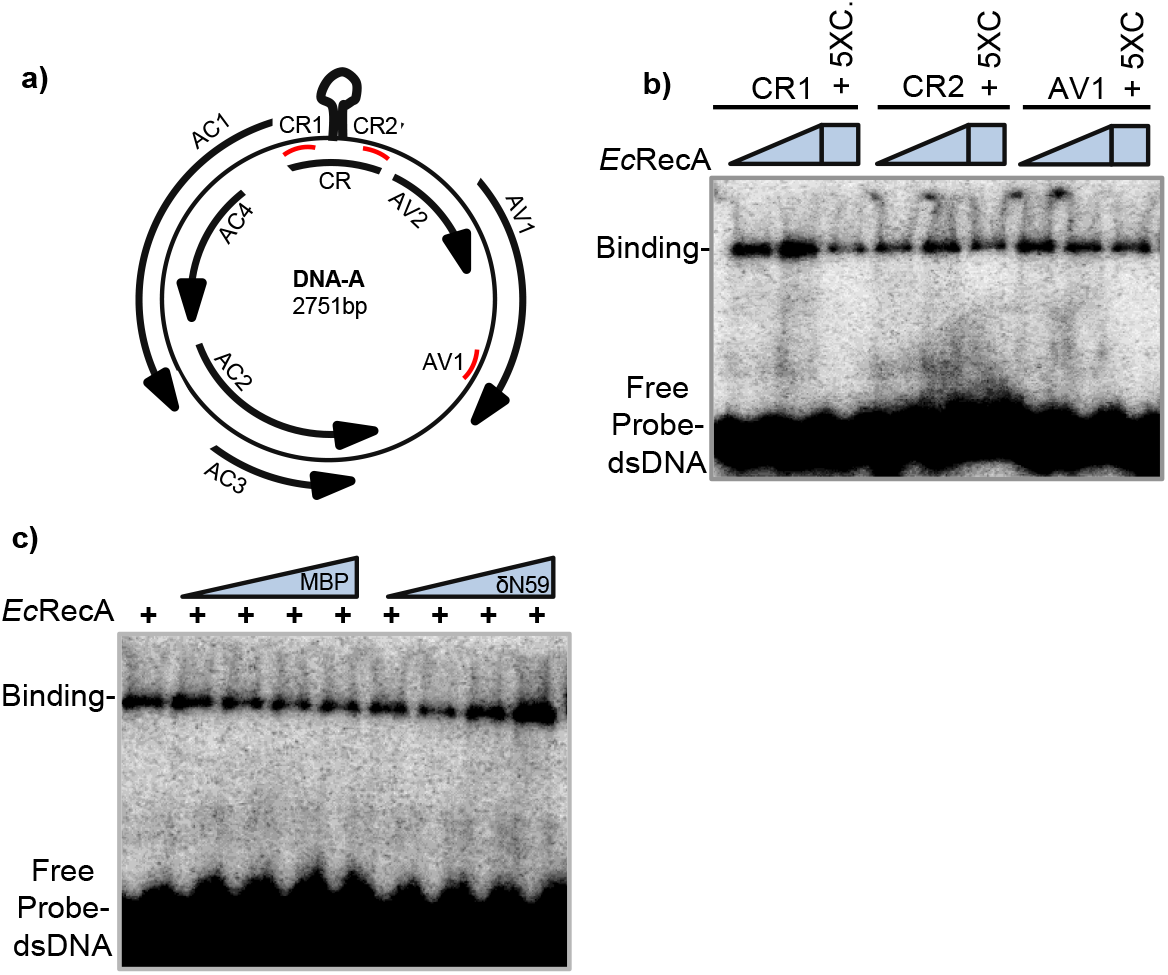
NtRecA1 binds to DNA. **a)** Schematic diagram showing SyYVCV genome architecture. Curved black arrow indicates ORFs and direction. Red line represents the region used to design probes. **b)** Interaction of *Ec*RecA with viral dsDNA probes. **c)** EMSA competition assay: RecA protein was co-incubated with viral DNA probe along with varied concentration of βC1 truncation mutant or MBP alone before resolving. 5XC indicates concentration of competitor (cold probe).

**Figure S8:**
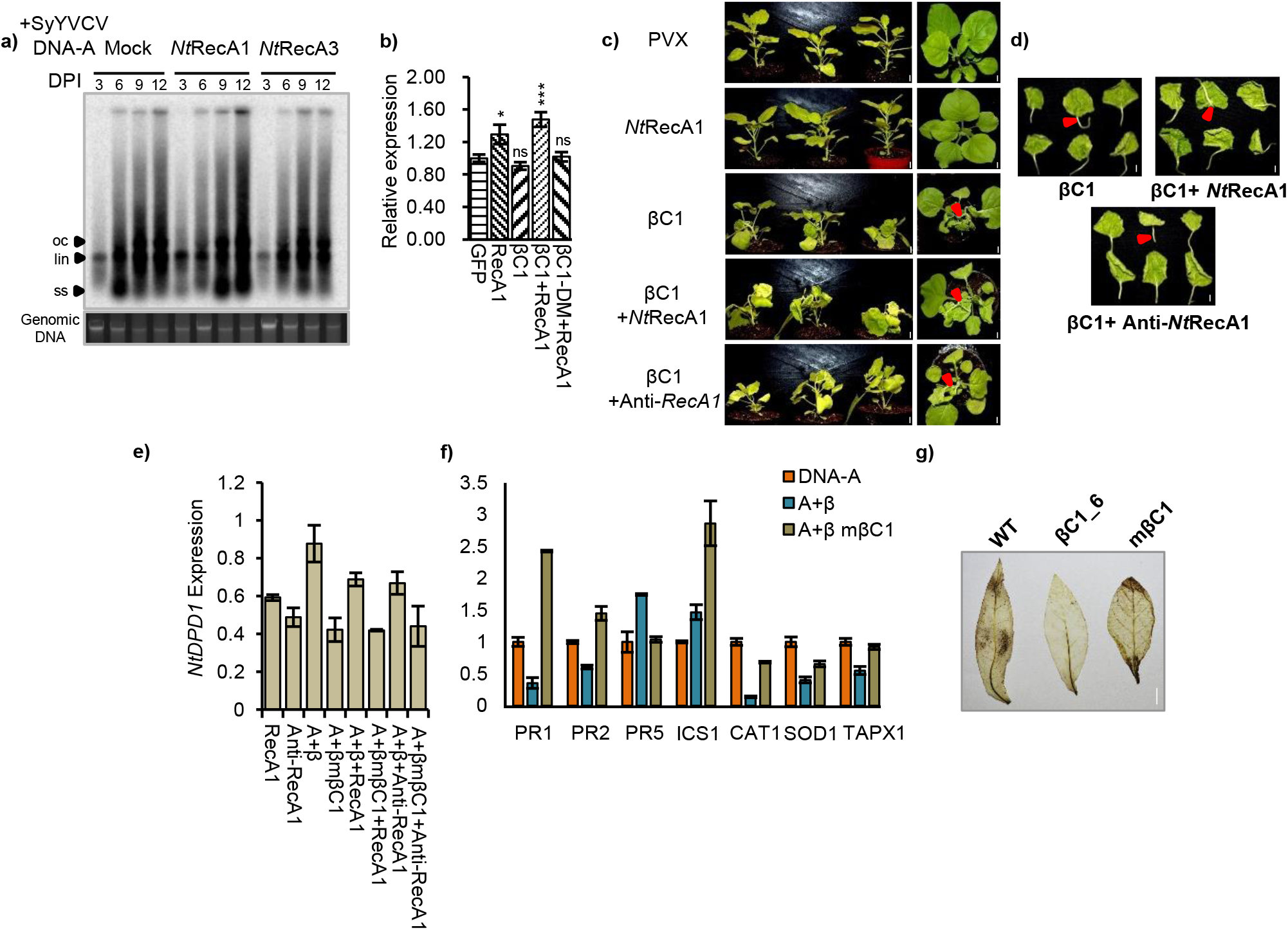
RecA1 enhances viral replication in the presence of βC1. **a)** Viral replication assay with SyYVCV DNA-A partial dimer co-inoculated with RecA1 or RecA3 in *N. tabacum*. SB blot was performed using full length DNA-A as probe. **b)** qPCR for viral titre analysis in *N. tabacum* infected with SyYVCV DNA-A in combination with βC1 or βC1-DM along with RecA1. **c)** RecA1 augments symptom determination function of βC1. Genes were expressed in PVX based vectors to analyse symptoms, either alone or in combinations. Images were taken at 12 DPI. *NtRecA1* or anti-*NtRecA1* expressed from PVX vector alone or in combination. **d)** Similar to c) except βC1 was used in combination with *NtRecA1* or anti-*NtRecA1*. Petiole was detached in each case. Red arrow indicates petiole angle. Scale bar 0.5 cm. **e)** Transcript expression of DPD1 during infection with various combinations. **f)** Transcript level of various PR and defense genes during infection with SyYVCV carrying WT βC1 or mβC1**. g)** DAB staining showing accumulation of ROS in βC1 and mβC1. Two week old tissue culture maintained transgenic plants were used for staining. Tukey’s multiple comparison test with three stars representing P-value, P ≤ 0.001 and two stars P ≤ 0.01. n=4.

**Figure S9:**
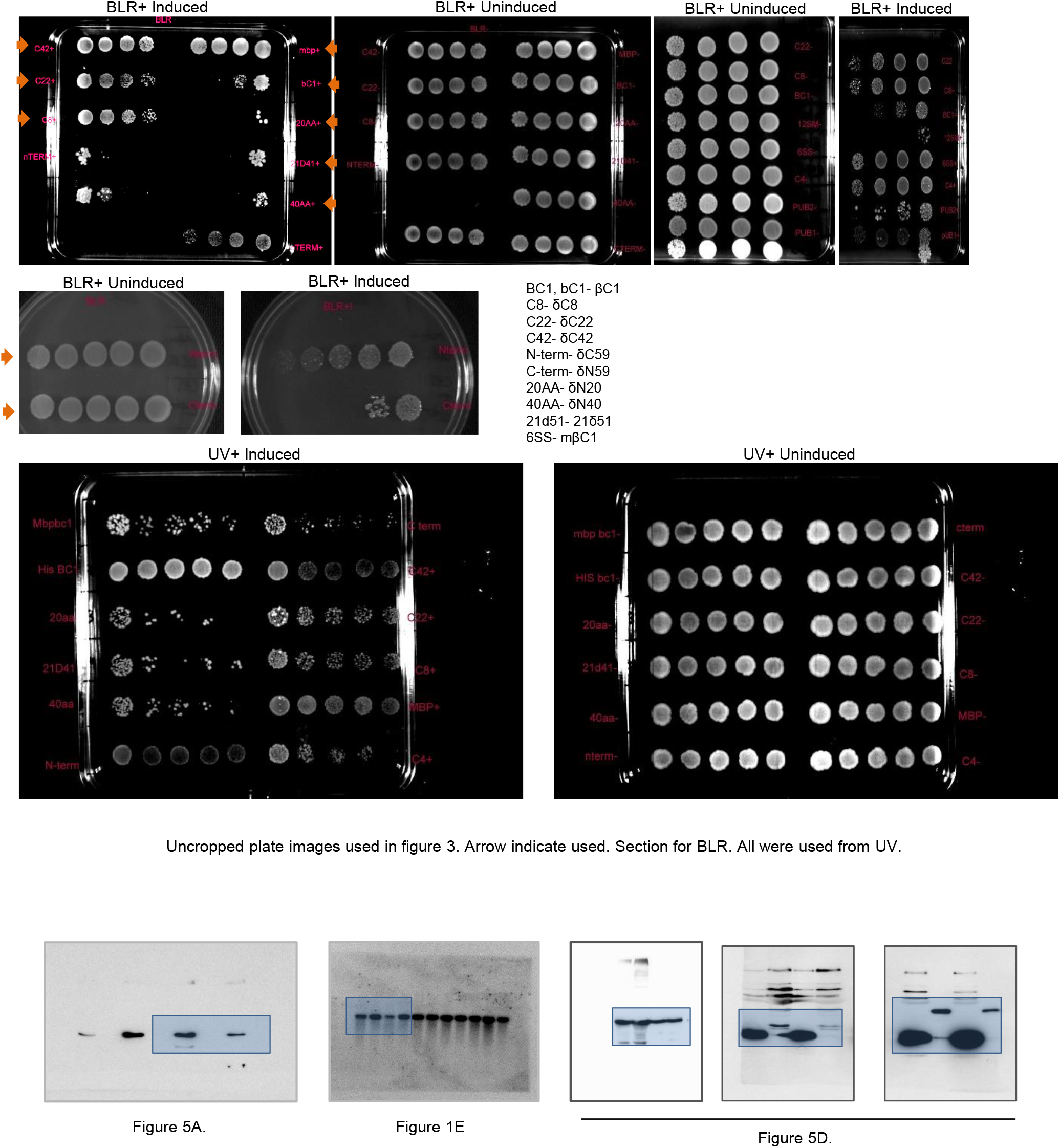
Full images of cropped blots and plates.

**Table S1:** Mass-spectrometry identified interacting proteins in *E.coli* purified βC1

| Accession | Description | Coverage [%] | Peptides | PSMs | Unique Peptides | Score Sequest HT: |
| --- | --- | --- | --- | --- | --- | --- |
| A0A0H2Z1Z1 | Protein RecA | 59 | 18 | 59 | 17 | 165.63 |
| A0A069XJD6 | Chaperone DnaK | 84 | 66 | 412 | 38 | 1366.38 |
| A0A0A0F6U9 | Chaperone DnaJ | 62 | 21 | 94 | 21 | 312.53 |
| A0A222QFX8 | Nuclease SbcCD | 43 | 30 | 43 | 4 | 124.39 |
| W8ZHA5 | Ribonuclease E | 56 | 44 | 68 | 8 | 202.58 |
| W8T8L9 | Exodeoxyribonuclease 7 | 20 | 6 | 21 | 6 | 19.24 |
| A0A166VD00 | Exoribonuclease 2 | 17 | 6 | 6 | 6 | 13.05 |
| A0A073FMW3 | RecF | 36 | 8 | 8 | 8 | 11.18 |
| A0A017IAN7 | RecBCD | 14 | 6 | 7 | 6 | 13.63 |
| A0A062Y8Q3 | RecN | 15 | 4 | 4 | 4 | 6.93 |
| A0A222QVR5 | RdgC | 23 | 4 | 4 | 4 | 2.18 |
| A0A1E5WYB1 | RecQ | 8 | 3 | 3 | 3 | 1.74 |

**Table S2:**
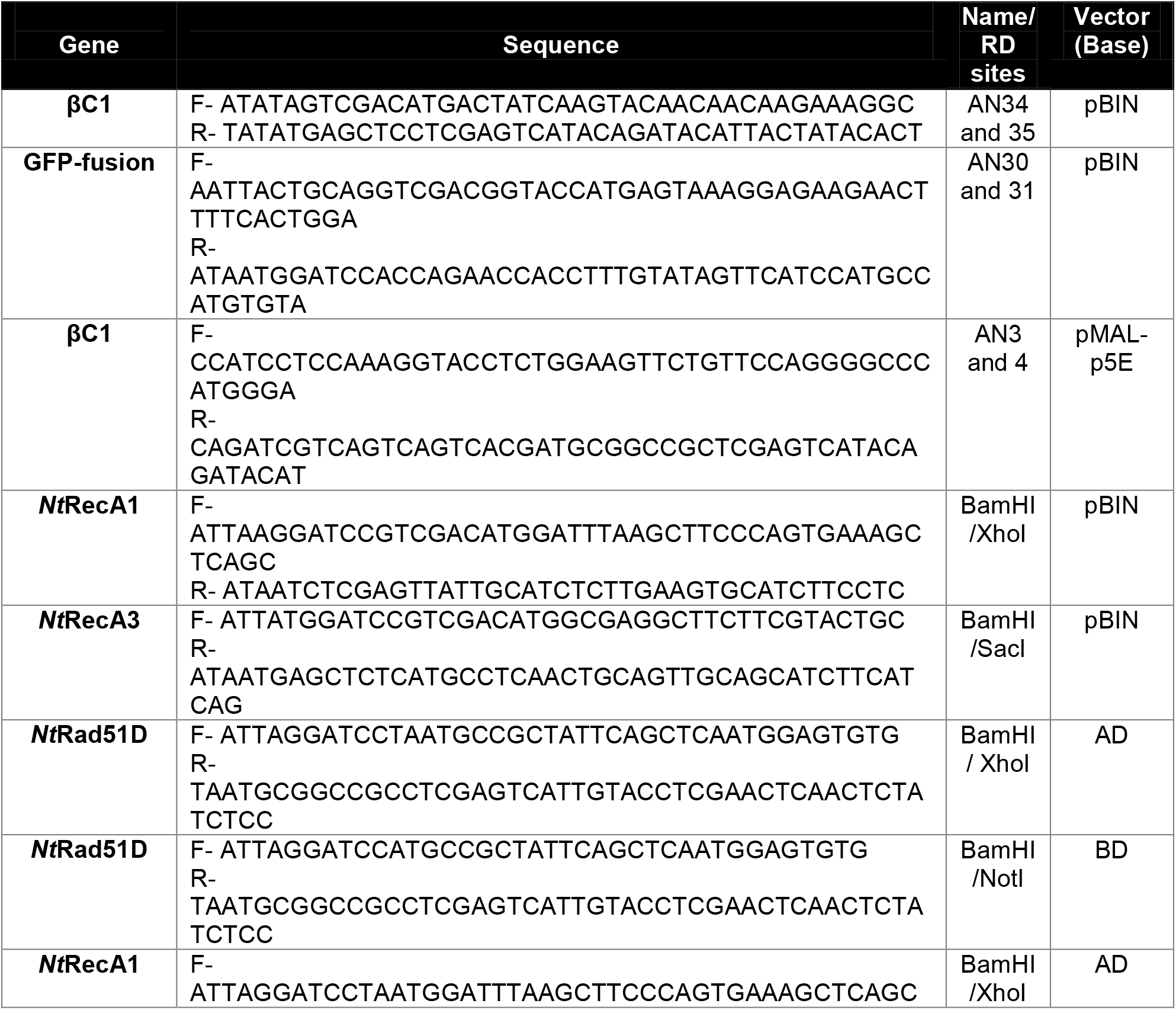
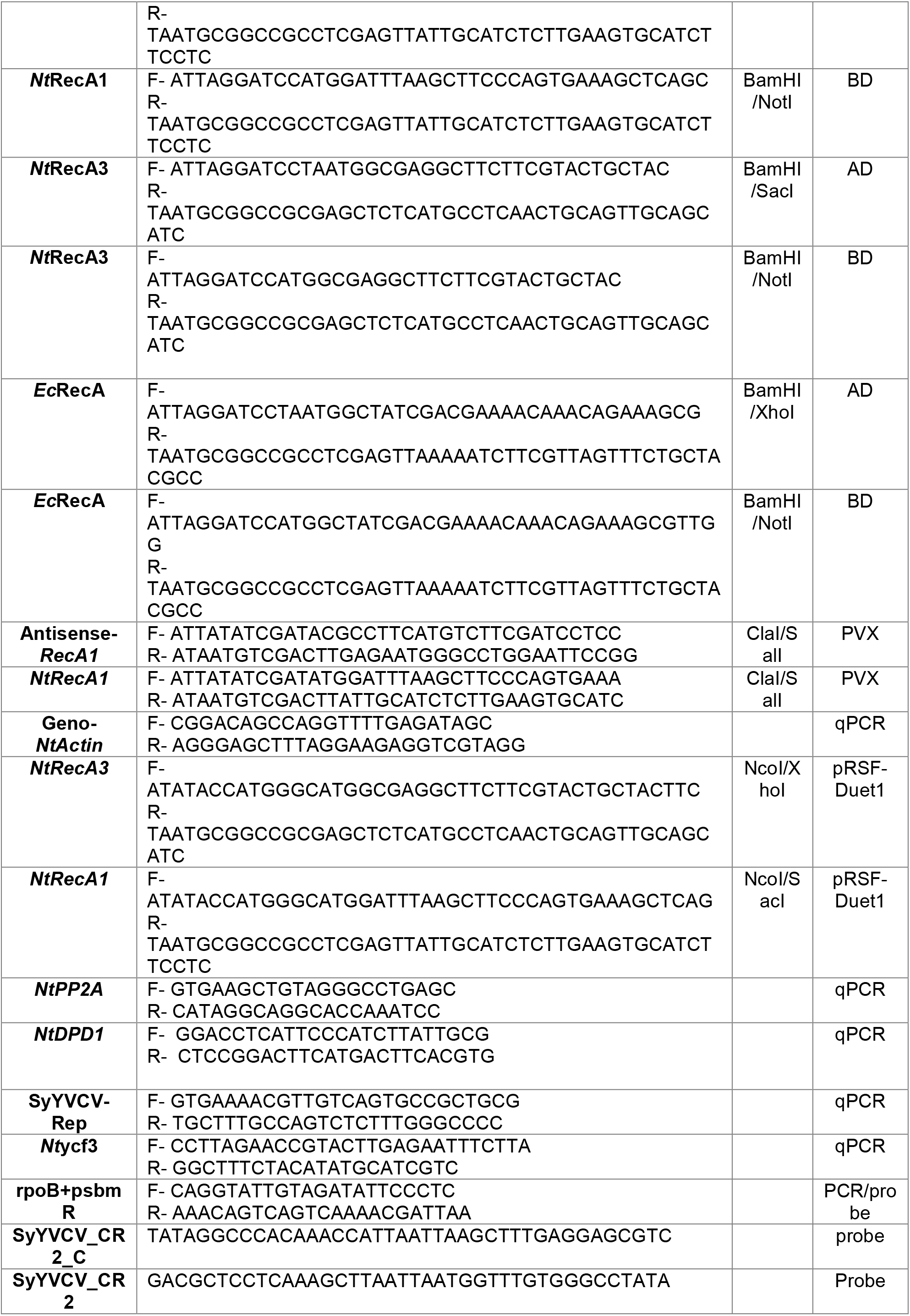
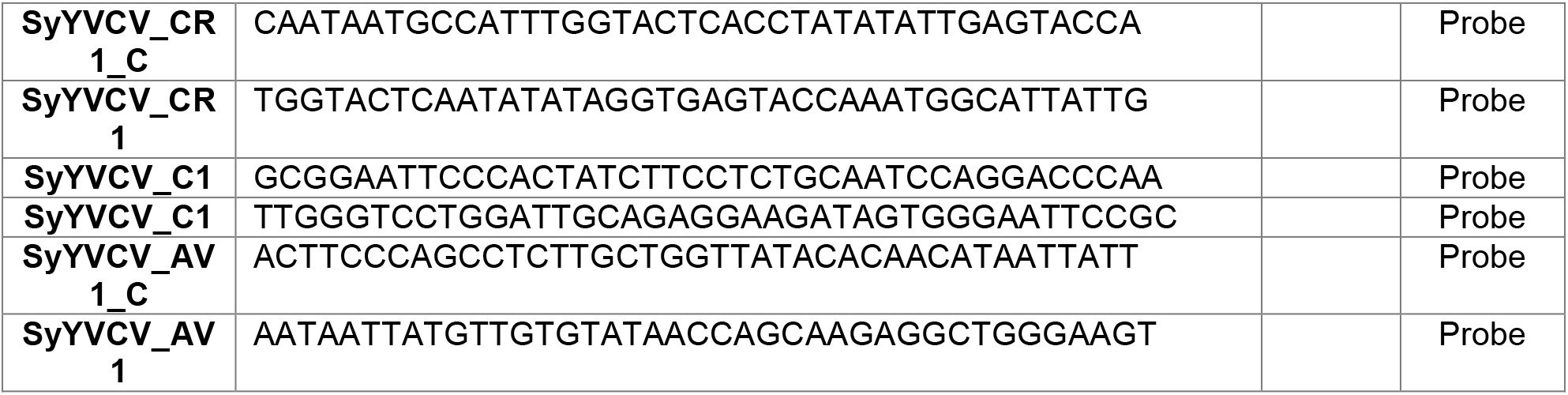
List of primers used in this study

**Table S3:** List of clones used in this study

| S.No | Gene name/ Tag details | Plasmid name | Vector (Base) |
| --- | --- | --- | --- |
| 1 | MBP-βC1 | pAN9 | pMAL-p5E |
| 2 | βC1-HIS | pAN17 | pBIN19 |
| 3 | βC1-GFP | pAN24 | pBIN19 |
| 4 | mβC1-GFP | pAN26 | pBIN19 |
| 5 | GFP-βC1 | pAN28 | pBIN19 |
| 6 | MBP-βC1_T8 | pAN31 | pMAL-p5E |
| 7 | MBP-βC1_m1 | pAN32 | pMAL-p5E |
| 8 | MBP-βC1_m2 | pAN33 | pMAL-p5E |
| 9 | MBP-βC1_m3 | pAN34 | pMAL-p5E |
| 10 | MBP-βC1_m4 | pAN35 | pMAL-p5E |
| 11 | MBP-βC1_m5 | pAN36 | pMAL-p5E |
| 12 | MBP-βC1_m6 | pAN37 | pMAL-p5E |
| 13 | MBP-βC1_m7 | pAN38 | pMAL-p5E |
| 14 | MBP-βC1_m8 | pAN39 | pMAL-p5E |
| 15 | MBP-βC1_m9 | pAN40 | pMAL-p5E |
| 16 | MBP-βC1_m10 | pAN41 | pMAL-p5E |
| 17 | MBP-βC1_T1 | pAN48 | pMAL-p5E |
| 18 | MBP-βC1_T5 | pAN49 | pMAL-p5E |
| 19 | MBP-βC1_T7 | pAN50 | pMAL-p5E |
| 20 | MBP-βC1_T6 | pAN51 | pMAL-p5E |
| 21 | MBP-βC1_T3 | pAN54 | pMAL-p5E |
| 22 | MBP-βC1_T2 | pAN55 | pMAL-p5E |
| 23 | MBP-βC1_T4 | pAN56 | pMAL-p5E |
| 24 | MBP mSIM2,3,4 βC1 | pAN78 | pMAL-p5E |
| 25 | AD-HA RecA | pAN79 | pGAD-T7 |
| 26 | AD-HA RecA1 | pAN80 | pGAD-T7 |
| 27 | AD-HA RecA3 | pAN81 | pGAD-T7 |
| 28 | BD-MYC Rad51D | pAN82 | pGBK-T7 |
| 29 | BD-MYC RecA | pAN83 | pGBK-T7 |
| 30 | BD-MYC RecA1 | pAN84 | pGBK-T7 |
| 31 | BD-MYC RecA3 | pAN85 | pGBK-T7 |
| 32 | DNA-A partial dimer | pSD-30 | pBIN19 |
| 33 | DNA-β partial dimer | AN-Beta& | pBIN19 |

**Table S4:**
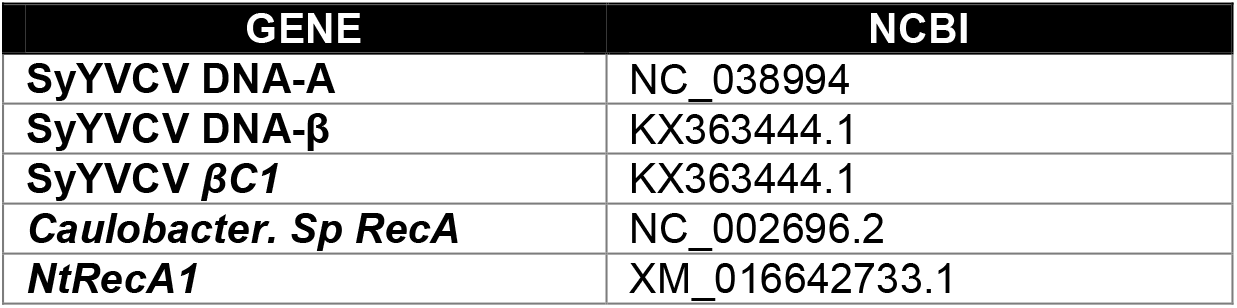
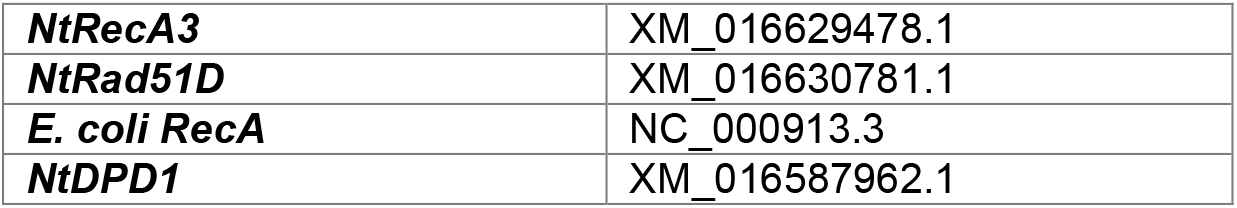
List of Sequence Id’s used to construct clones

**Table S5:** List of antibodies and IP materials used in this study

| **Reagent Type** | **Material** | **Catalogue No.** |
| --- | --- | --- |
| Antibody | Anti-GFP | Abcam (ab290) |
| Antibody | Anti-MBP | Abcam (ab9084) |
| Antibody | Anti-His | CST (2366) |
| Antibody | Anti-MYC | Abcam (ab9106) |
| Antibody | Anti-HA | CST (3724) |
| IP | GFP-Trap | Chromotek |
| IP | Anti-MBP-Magnetic | NEB (E8037S) |
| Beads | MBP(Dextrin Sepharose) | GE (28935597) |
| Beads | Ni NTA (Agarose) | Qiagen 30210 |

